# CoralBlox: A computationally efficient coral model for decision support

**DOI:** 10.64898/2026.04.13.718315

**Authors:** Pedro Ribeiro de Almeida, Rose Crocker, Daniel Tan, Kevin R. Bairos-Novak, Chinenye J. Ani, Jessica A. Benthuysen, Barbara J. Robson, Samuel Matthews, Takuya Iwanaga

## Abstract

Coral reef management under climate change is challenging due to data sparsity and high uncertainty, yet it is essential for informing conservation strategies. We present CoralBlox, a mechanistic discrete time coral ecology model with the explicit aim of supporting rapid scenario exploration and decision making. The model represents discretized distributions of five coral functional groups across configurable spatial scales while incorporating key ecological processes, including coral growth, reproduction, thermal adaptation, and responses to disturbances. Validation against observed data demonstrates that CoralBlox effectively captures major trends in coral cover dynamics across the Great Barrier Reef, particularly for bleaching-driven mortality and recovery patterns. While simplifying ecological complexities, the model maintains sufficient ecological realism to evaluate and compare the result of distinct management strategies. CoralBlox enables comprehensive assessment of potential management interventions with high computational efficiency and interoperability. The model’s flexible architecture makes it extensible to coral ecosystems worldwide, providing valuable exploratory capability for reef management.

**Teaser:** CoralBlox is an efficient coral reef ecology model supporting rapid scenario testing and management decision making under climate change.

**Highlights:**

- Marine ecosystems are characterized by high uncertainty and data sparsity.
- Management decisions still need to be made under these uncertain contexts.
- CoralBlox offers a conceptually simple yet credible representation of ecological processes.
- Comparatively fast runtimes across different spatial scales enable rapid exploration of plausible future states.

## 1. Introduction

Coral reefs are increasingly impacted by climate change and are projected to continue to decline without mitigation efforts (1). Effective coral ecosystem management requires considering multiple plausible futures across diverse potential strategies. Decisions must be made in a timely and informed manner with consideration of known risks and uncertainties, requiring the ability to quickly and robustly assess decision-relevant outcomes, identify acceptable alternatives, and discover new effective management strategies under *deep uncertainty* (2–4).

The GBR encompasses multitudes of ecological and biological processes that are influenced by global oceanographic and climatic processes (5) alongside anthropogenic impacts that negatively impact reef ecosystems. While comprehensive reef monitoring does exist – in the form of the Australian Institute of Marine Science (AIMS) Long-Term Monitoring Program (6) and the Great Barrier Reef Marine Monitoring Program (7), there remains a sparsity of longitudinal demographic data to support long-term projections of ecosystem responses under plausible future environmental conditions. Available long-term monitoring data covers approximately eight percent of reefs recognized by the Great Barrier Reef Marine Park Authority (296 of 3,806) (8) across 344,400 km^2^.

The combination of complexity, data sparsity, parameter uncertainty and lack of consensus characterizes the situation as one of *deep uncertainty* (4), wherein decision-makers must act without shared understanding of key drivers of change or their potential evolution over time. Disagreement on how a system works, which trends matter most, or how probable different future pathways may be is a common feature of deep uncertainty. Further examples of uncertainty include the vast array of environmental and climatic futures (such as the magnitude and extent of any coral bleaching and cyclones), the shape and magnitude of restoration interventions and their expected efficacy (9–12).

Models that are increasingly complex and include many additional subprocesses that are not well understood may reduce overall model performance, reduce model transferability/adaptability, and experience reduced uptake by managers. The inclusion of more uncertain parameters or variable processes can increase overall model uncertainty, especially when data is sparse (13). Additionally, increasing model complexity can reduce the generality of models and thus their transferability and adaptability to novel conditions (14), which may also reduce model uptake by managers (13). Finally, increasingly complex models are more difficult to communicate to managers, reducing their uptake in decision-making processes (13). Thus, a balance of including only key known, well-studied processes and associated parameters that are in accordance with the scale of decision making is crucial.

Nevertheless, decision support remains valuable and becomes even more important under *deep uncertainty*. Although individual projections remain unreliable, the uncertainty in outcomes attributable to different management strategies can be significantly smaller than the models’ broader predictive uncertainty (15–17). The aim is to identify management strategies that robustly achieve acceptable outcomes (i.e. over a vast array of plausible futures), understand the conditions under which they fail, and to provide indications of when managers should consider changing tack (18–20). This study proposes a novel framework to explore a wide range of plausible future conditions such that the emphasis shifts from optimality and precise prediction to flexibility and meaningful comparisons between management options. Such large-scale scenario exploration, particularly in the context of a future operational decision support, demands computationally efficient models.

CoralBlox (21) is a computationally efficient coral ecosystem model for the Great Barrier Reef (GBR) with runtimes in the order of seconds. It is designed to explore the regional implications of management decisions for the GBR. CoralBlox is a companion model bundled with the Adaptive Dynamic Reef Intervention Algorithms (ADRIA) decision support platform (22). Although CoralBlox has been designed for decision support within the GBR as its primary use case, the model can be applied to any coral reef ecosystem, provided sufficient data are available.

CoralBlox addresses three critical needs: 1) rapid and iterative analyses for exploratory purposes; 2) parsimony combined with conceptual defensibility and scientific rigor through demonstrated ability to reproduce past observations; and 3) support for iterative stakeholder discussions where comparatively short timeframes between consultation and feedback is preferable. CoralBlox achieves these requirements by representing the coral ecosystem through a simplified model structure and parameterization that retains key elements of the dynamic system. Model parsimony has the additional benefit of having reduced parametric uncertainty, easing subsequent analysis and assessment (23).

The central question addressed in this paper is: can such a simplified, size-structured representation of coral ecosystem dynamics reproduce observed coral cover trajectories on the GBR with sufficient skill to support management decision-making? We address this by (1) testing CoralBlox’s ability to reproduce historical coral cover trajectories against independent calibration and test reef sets; (2) quantifying model performance at both individual-reef and regional scales; and (3) using global sensitivity analysis to identify which ecological and environmental factors most strongly influence model outcomes. Together, these results establish the conditions under which CoralBlox’s efficiency-for-fidelity trade-off is justified, and outline future use cases for CoralBlox in decision support on the GBR and globally.

## 2. Results

### 2.1 Model overview and parameterization

CoralBlox is a coral ecosystem dynamics model designed to enable the rapid exploration of many possible futures for the Great Barrier Reef. At discrete (annual) timesteps, the model tracks coral size (diameter) distributions for five functional groups discretized into cohorts referred to as *blocks*. The range of allowed sizes for each functional group is split in seven size classes, so that corals within the same size class and functional group share the same growth and mortality rates. Although the size classes remain constant for the entire simulation, at each timestep these *blocks* move within and between size classes and “shrink” in height to account for growth and mortality, respectively (Figure 1B). The “Blox” portion of the model’s name refers to this methodological approach.

**Figure 1.**
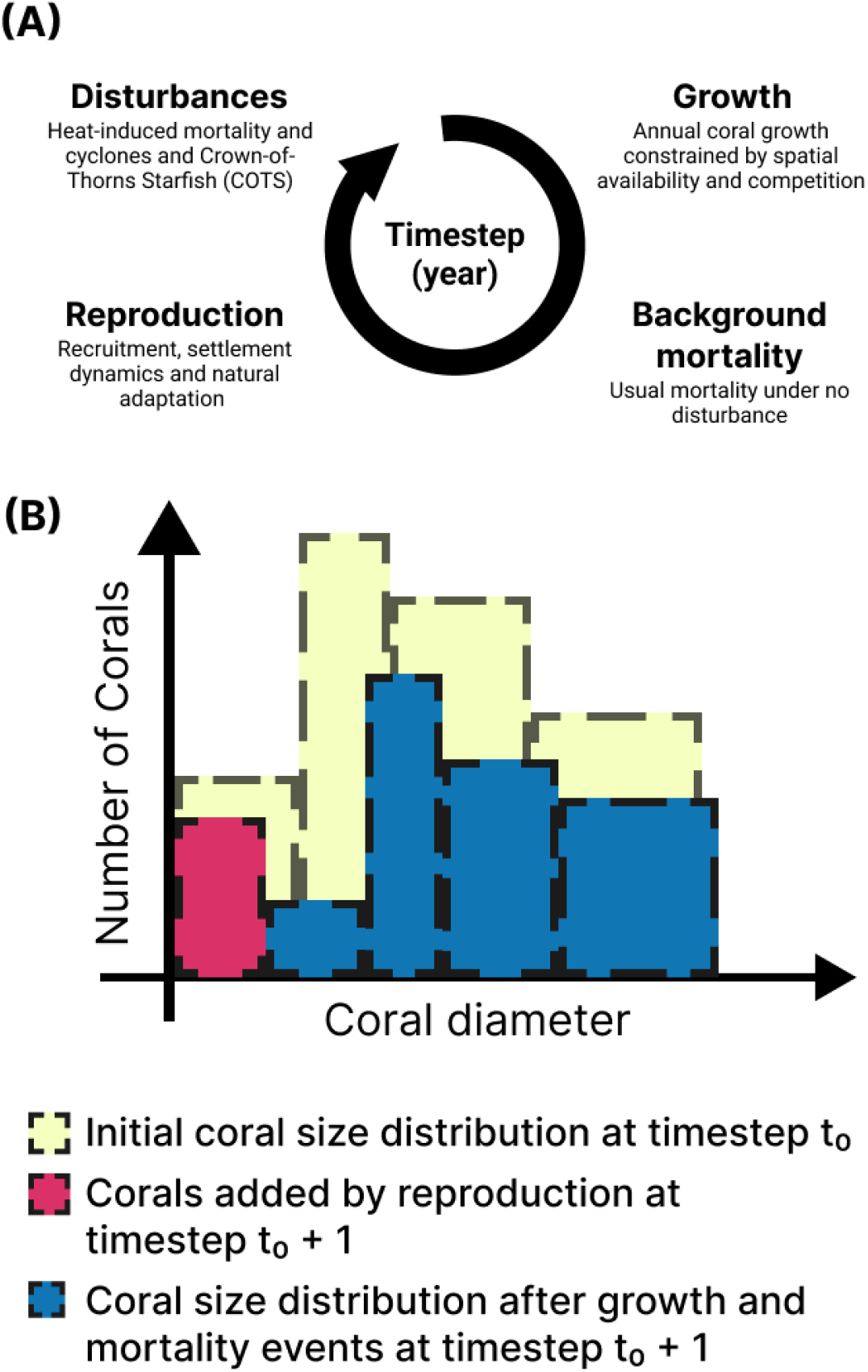
Ecological and environmental processes represented by CoralBlox within a timestep. (A) Sequence of five processes that take place, in that order, over a timestep in CoralBlox for each reef and each functional group. Coral mortality caused by COTS is not modelled but is represented via forcing data. (B) Illustration of the discretized size distributions before (yellow) and after (magenta and blue) a timestep. Each distribution is composed of chunks of discretized diameter distribution (*blocks*) of different heights and widths that, together, represent the diversity of coral diameters in the population. The horizontal movement and shrinkage of each block represent coral growth and mortality. The magenta block accounts for new coral entering the system due to reproduction.

CoralBlox is a network-based meta-population model that is spatially structured but not spatially explicit over a reef environment. Any arbitrary area where corals may live is characterized by its physical properties (median depth, total spatial area, and coral habitable area) and can represent any nominal spatial scale (i.e. whole of reef or sub reef). Larval connectivity is represented through a transfer probability matrix that provides the likelihood of larvae moving between represented locations. Distinct ecological and environmental processes that align with ecological lifecycles are represented sequentially on an annual basis. These processes include growth, background mortality, reproduction and response to environmental disturbances owing to climatic events or biological threats (i.e. bleaching mortality, cyclone damage and predation by Crown-of-thorns Seastar (COTS) (Figure 1). Model parameterization enables representation of these ecological and environmental processes as functions of the functional groups and size classes (explained further in Materials and Methods). A single CoralBlox simulation’s computational cost (3806 reefs over 15 years) was 1.3 ± 0.1 seconds (mean ± standard deviation over 10 repetitions, single-core and single-threaded, on a standard laptop).

### 2.2 Reproducing coral cover trajectories on the GBR: 2008-2024

Only a fraction of the reefs monitored by the LTMP have enough observations to support a model performance assessment. We split these into a 75% calibration set and a 25% test set: model parameters were fit to the calibration set, while the test set was reserved to assess performance on unseen reefs. Refer to the “Model calibration and testing” section of the Materials and Methods for further details.

Figure 2 presents a spatial overview of the performance metrics used to assess the model: the difference between modelled and benchmark Root Mean Square Errors (Δ*RMSE*) and the Spearman Rank Correlation Coefficient (SRCC). These were calculated individually for each reef in the calibration and test sets. Table S1 presents the bootstrapped median RMSE, ΔRMSE, SRCC, median bias and median absolute bias for calibration and test reefs, with 95% confidence intervals. The full distribution of scores underpinning Figure 2 and Table S1 is presented in Figures S5-S8.

**Figure 2.**
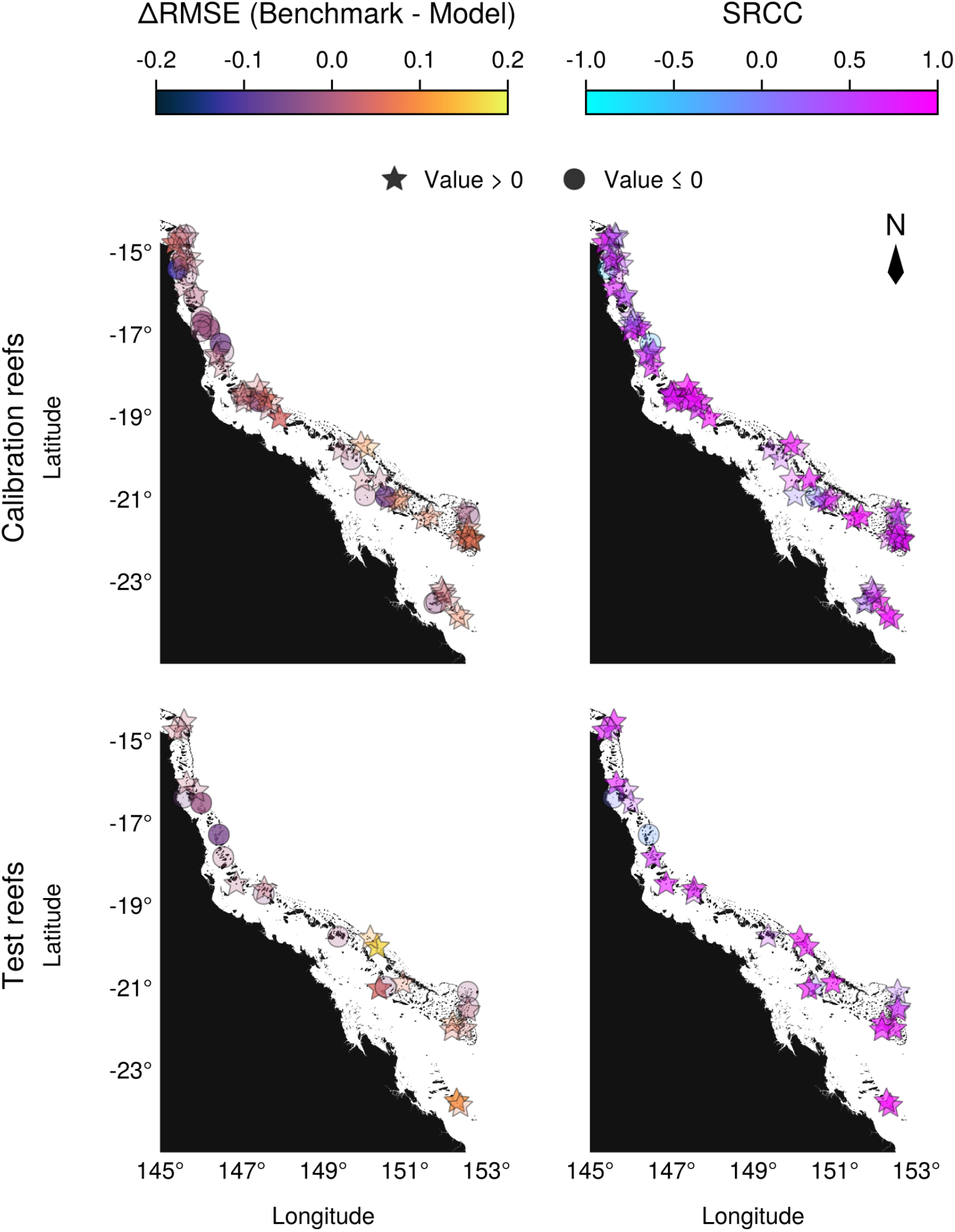
Model performance metrics for calibration and test reefs. Positive values are represented by stars and negative values by circles. First column: difference between historic average and model’s RMSE (**ΔRMSE**) for each reef. Positive values indicate the model outperformed the benchmark. The model outperformed the benchmark at 50 of 73 reefs (for calibration) and 17 of 26 reefs (for test). Second column: Spearman’s Rank Correlation Coefficient (SRCC) between model and observation for each reef. Bootstrapped median values with confidence intervals are: for calibration reefs **ΔRMSE**, 0.02 [0.01, 0.04], for test reefs **ΔRMSE**, 0.02 [-0.0, 0.05], for calibration reefs **SRCC**, 0.77 [0.7, 0.82], and for test reefs SRCC, 0.82 [0.55, 0.89].

Table S2 presents the spatial and heat-stress correlation coefficients (SRCC) with respect to each reef’s performance metric. Positive spatial correlations both along and across the GBR indicate an increase in the performance metric southward and offshore, respectively. For heat-stress correlations, negative correlations indicate a decrease in the performance metrics with increasing DHW.

Figures 3 and 4 present the mean relative cover from 2008 to 2022 across each *reef group* (Figure S1) for calibration and test data splits, respectively. The median RMSE and SRCC for each *reef group* is also indicated. Unlike the other performance metrics reported in this study, these reef-group-level trends are not assessed via bootstrap resampling since most reef groups have too few reefs for meaningful resampling (as few as two reefs in some test reef groups). In both figures, consecutive years of the observed trajectory are connected by a solid line, so dots that appear unconnected on one or both sides indicate a gap in the historical data. The shaded intervals in these figures show the 95% range of cover across the reefs within each group.

**Figure 3.**
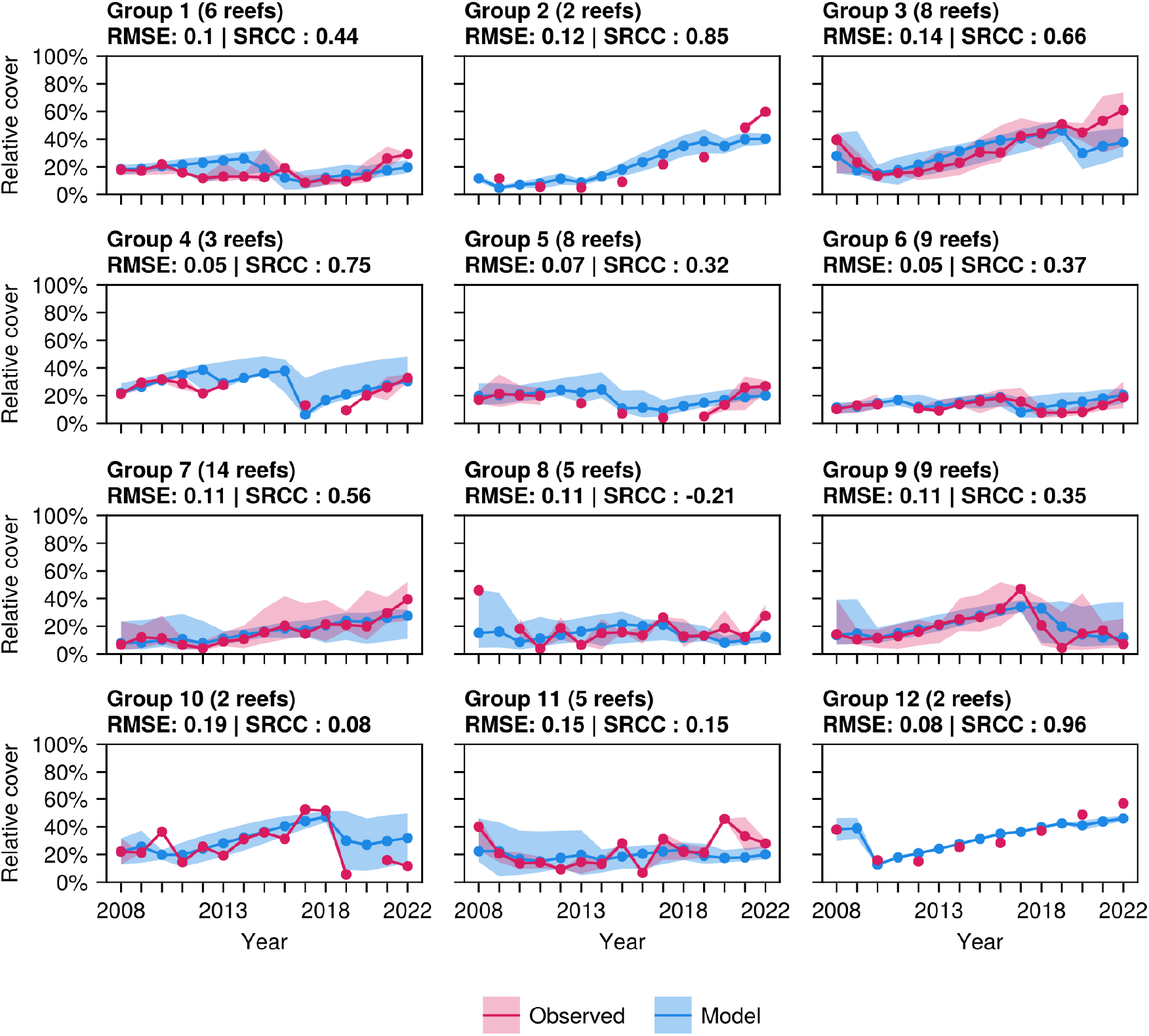
Observed versus modelled relative coral cover for calibration reefs aggregated by reef group. Shaded intervals show the 2.5th-97.5th percentile range of cover across reefs within each group.

**Figure 4.**
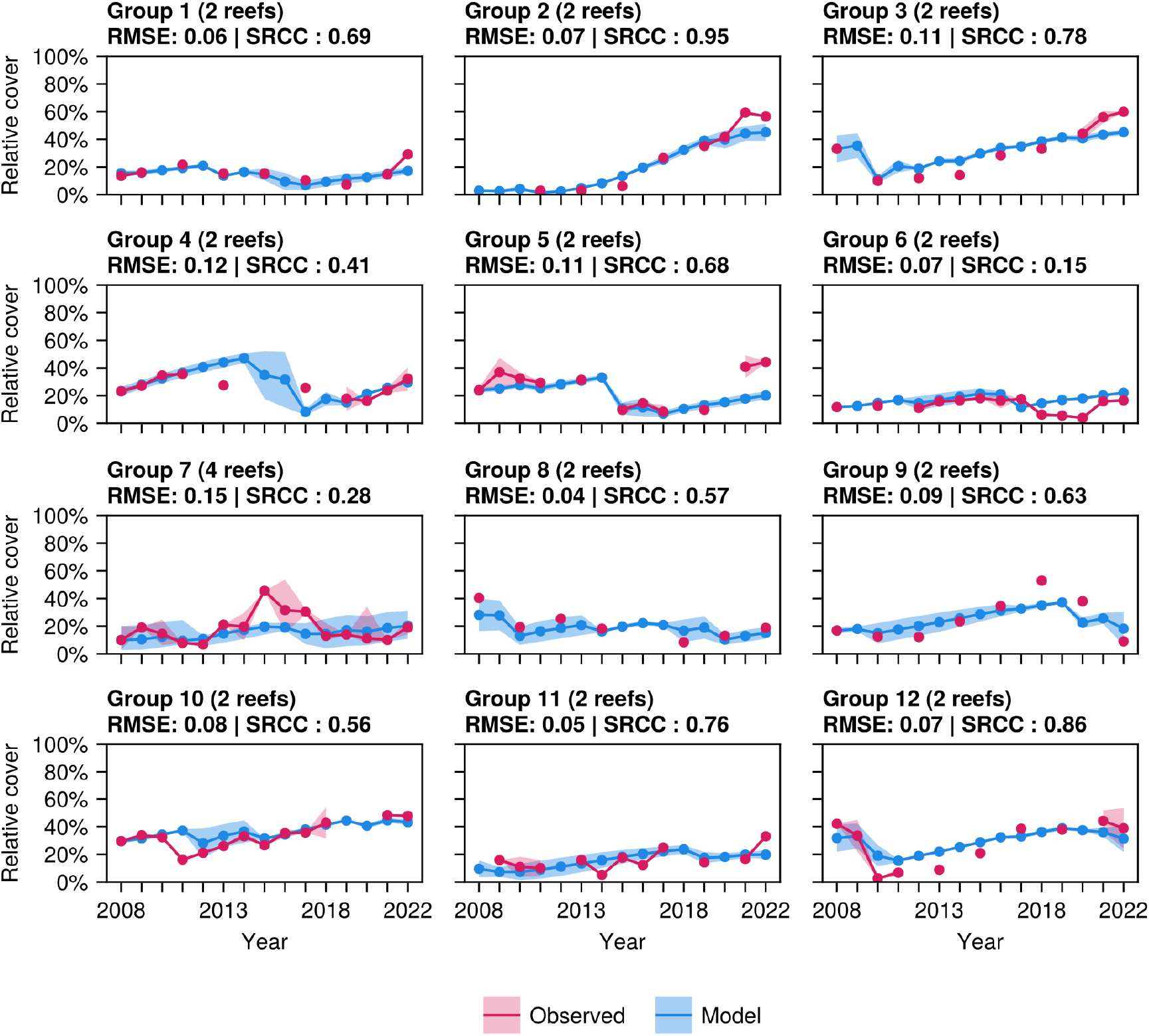
Observed versus modelled relative coral cover for test reefs aggregated by reef group. Shaded intervals show the 2.5th-97.5th percentile range of cover across reefs within each group.

A comparison between modeled and observed relative coral cover for four selected test reefs is presented in Figure 5. The selected reefs (the two with the lowest and two with the highest ΔRMSE) show a range of distinct environmental and ecological behaviors, including low and high heat stress, presence or absence of mortality events caused by cyclone or Crown-of-thorns Starfish (COTS) outbreaks, low and high initial coral cover, and distinct initial benthic compositions. Cyclone and storm damage historical data and the lack of a COTS sub-model make validating model processes difficult at reefs significantly impacted by these disturbances. Estimates of mortality for these events are inferred directly from the LTMP to ensure trajectories are aligned.

**Figure 5.**
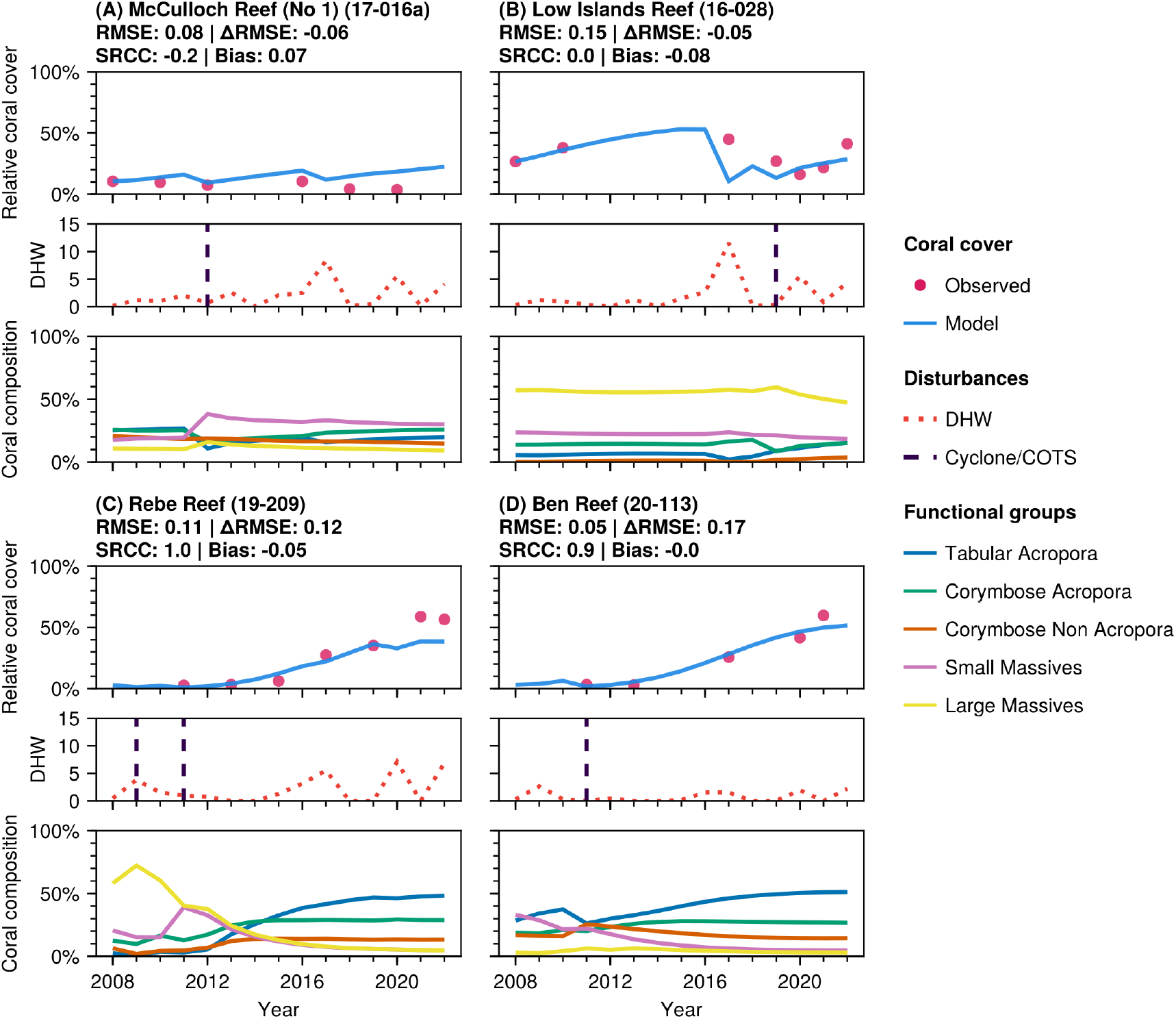
Comparison between modelled and observed coral cover for selected reefs. (**A**) **McCulloch Reef (No 1) (17-016a):** Model RMSE = 0.08, Benchmark RMSE = 0.03, ΔRMSE = −0.06, SRCC = −0.2, Bias = 0.07; (**B**) **Low Islands Reef (16-028):** Model RMSE = 0.15, Benchmark RMSE = 0.1, ΔRMSE = −0.05, SRCC = 0.0, Bias = −0.08; (**C**) **Rebe Reef (19-209):** Model RMSE = 0.11, Benchmark RMSE = 0.23, ΔRMSE = 0.12, SRCC = 1.0, Bias = −0.05; (**D**) **Ben Reef (20-113):** Model RMSE = 0.05, Benchmark RMSE = 0.22, ΔRMSE = 0.17, SRCC = 0.9, Bias = −0.0. For each reef, the subplots show the modelled (solid blue line) and observed (red dots) values of relative coral cover; the historical DHW values (red dotted line) and historical mortality events caused by Crown of Thorns Starfish (COTS) and/or cyclone damage (vertical dashed lines); and the modelled relative coral composition.

### 2.3 Coral survival under high DHW varies with depth

We performed a variance-based global sensitivity analysis using Shapley effects (24, 25) to determine how distinct ecological and environmental factors influence four metrics: change in coral cover, shelter volume (measured in *m*^3^ (26)), evenness (measure as the inverse of the Simpson’s Index (27)) and juvenile cover (measured as the coral cover for the first two smaller size classes) over a single simulated timestep (Figure 6). The implementation of these metrics is described in (28). To produce each plot in Figure 6, we used 30 000 samples per DHW range. See Figure S4 for a convergence analysis. A synthetic (hypothetical) reef was used for this analysis, enabling parameterization of some of the reef’s environmental and ecological features (namely, habitable area, depth, and benthic composition); initial relative cover itself was held fixed at 0.3 across all scenarios, isolating the sensitivity analysis to compositional and environmental effects rather than absolute starting cover. Refer to Table S11 for a full list of all factors included in this analysis (219 in total).

**Figure 6.**
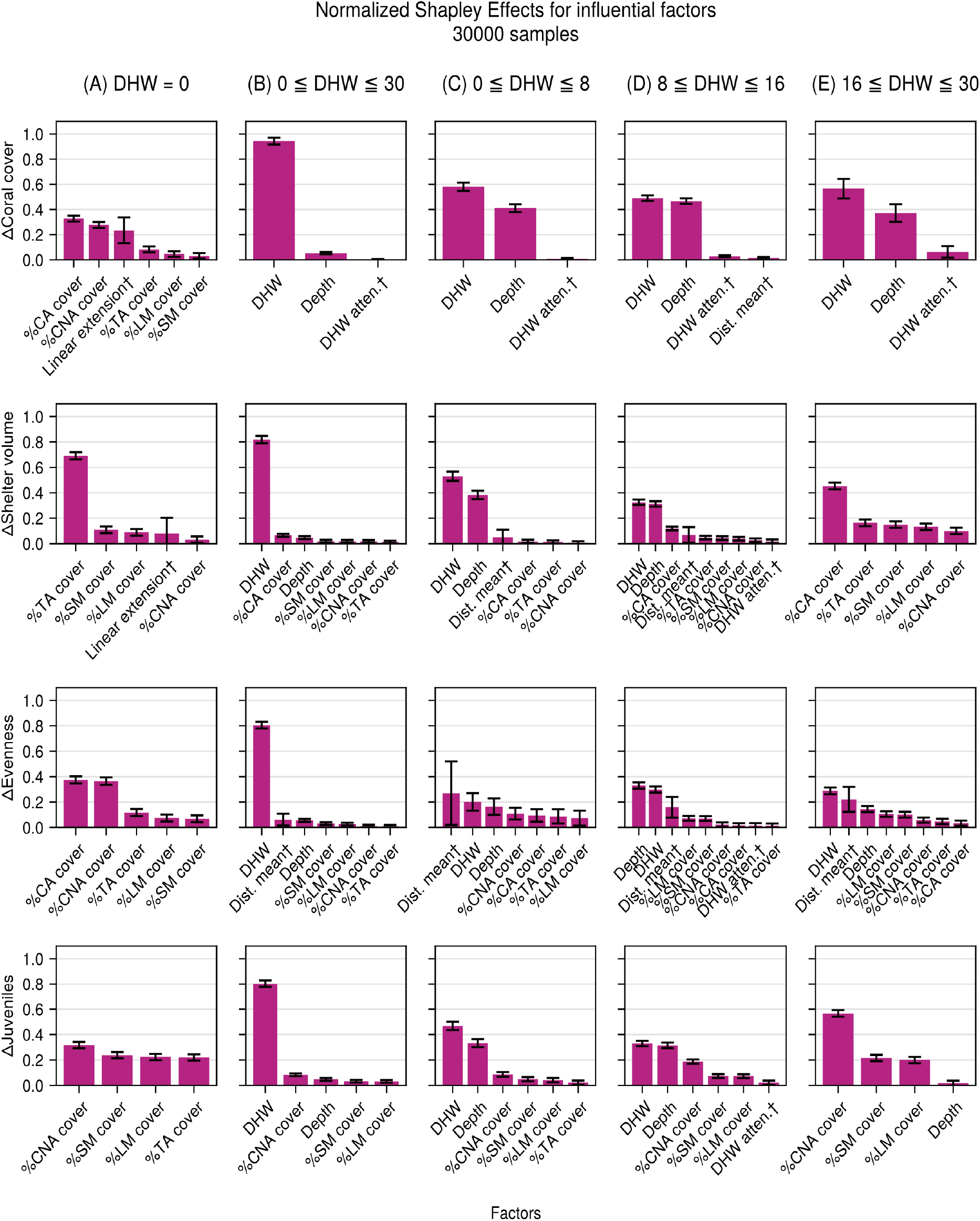
Shapley effects for distinct disturbance scenarios. Factors with a † represent groups of parameters across distinct functional groups, size classes or reef groups. Initial proportion of each functional group – tabular Acropora (%TA), corymbose Acropora %CA, corymbose non-Acropora (%CNA), small massive (%SM), and large massive (%LM) – were included in the assessment. (A) DHW = 0: no heat stress scenarios. (B) 0 ≤ DHW ≤ 30: Broad-range heat stress scenarios. (C) 0 ≤ DHW ≤ 8: Low-value and narrow-range heat stress scenarios. (D) 8 ≤ DHW ≤ 16: Mid-value and narrow-range heat stress scenarios. (E) 16 ≤ DHW ≤ 30: High-value and narrow-range heat stress scenarios.

We included a dummy factor – a model input known to have no effect on model outcomes – to provide a statistical threshold (29). From the 219 factors included in the analysis, only the ones whose sensitivity indexes were higher than the dummy factor were included in the analysis. Under no heat stress (Figure 6A), linear extension and benthic composition (in particular for corymbose Acropora and corymbose non-Acropora) appear as the most influential factors for change in coral cover. In scenarios with a broad range of heat stress (Figure 6B), DHW dominates change in coral cover. Stratifying heat stress into low, medium and high ranges (Figure 6C**-**E) shows that, for this same metric, depth’s influence rivals DHW’s only in the medium-heat-stress range, where the depth-refuge effect can be the difference between a reef surviving or not. At low DHW, heat stress is rarely severe enough for depth to matter; at high DHW, it is severe enough to overwhelm the protection depth provides regardless of location. Depth’s contribution to coral cover therefore peaks in this intermediate heat-stress window, where its buffering capacity is neither redundant nor overwhelmed.

## 3. Discussion

CoralBlox provides a computationally efficient modelling capability to support rapid screening of potential management approaches. By explicitly representing key ecological processes within a simplified but credible framework, CoralBlox enables stakeholders and reef managers to iteratively explore management options and quickly identify robust strategies under deep uncertainty. CoralBlox balances the need for credible representations of ecological interactions (at reef-regional scales) and computational efficiency to provide timely, actionable insight. This is reflected in the bias metrics reported in Table S1: CoralBlox is intended to give a good indication of regional coral cover dynamics rather than to precisely predict any individual reef’s trajectory, so the near-zero signed median bias is relevant at the regional scale, while the larger median absolute bias reflects the expected margin of error at the individual reef scale.

The comprehensive approach taken to calibration and testing provides key insights to the development of coral ecosystem models and intervention management on the GBR. Model performance metrics increased southward and offshore and decreased with increasing DHW exposure. Furthermore, the model displayed highest sensitivity to DHW and depth (Figure 6). Together these findings indicate that the increased influence of bleaching events since 2016 increases uncertainty in model projections. This adds to mounting evidence that DHW alone cannot fully resolve coral bleaching mortality events (30) and that regional ecosystem models should account for factors such as cloud cover (i.e. light attenuation) (31), turbidity (32) and an array of other modifying variables (33). Indeed the decrease in model performance in inshore reefs is consistent with other GBR regional models and indicative of the mediating effect that water quality and turbidity play in bleaching mortality and recovery rates (34). This suggests that future projections of the GBR (35, 36) need to be interpreted carefully as ameliorating factors for the most sensitive components of these models (i.e. related to bleaching mortality) are not routinely incorporated. Importantly however, these results support recent findings indicating that management of deeper reefs may play a crucial role in in the future of the GBR with increased exposure to severe bleaching events (37).

Balancing model complexity with performance allows the simulation of thousands of potential management scenarios, enabling stakeholders to better explore intervention options and their implications. While the system is currently limited in its representation of non-heat related disturbances such as COTS outbreaks, validation against historical data shows that CoralBlox can effectively capture bleaching impacts and recovery dynamics in many reef contexts. As a complementary component of the Reef Restoration and Adaptation Program’s modelling toolkit (35, 38), CoralBlox provides the first computationally efficient platform to develop informed and timely interventions to enhance coral reef resilience in the face of increasing climate pressures.

### 3.1 Limitations and future work

While CoralBlox provides a framework for rapid reef management scenario exploration, it comes with several limitations. The model employs simplified representations of complex ecological processes and interactions that inevitably sacrifice some mechanistic details and accuracy for computational efficiency. Spatial heterogeneity within coral reef environments is represented at a coarser scale than in more complex models, potentially overlooking fine-scale ecological dynamics that may influence outcomes in certain contexts. CoralBlox is first and foremost intended for exploratory scenario analysis to support decision-making processes.

CoralBlox simulates a set of functional groups similar to those used in ReefMod-GBR (39) and C~scape (38). The decision was made in part to ease communication with reef managers and other reef scientists by adopting familiar and tested ecological representations to explore decision options and outcomes. The current implementation does not yet fully represent several important disturbances, most notably COTS outbreaks, cyclones and water quality impacts. As demonstrated in our validation against historical observations, this can lead to discrepancies in coral cover trajectories at reefs strongly affected by these disturbances. Not representing COTS in projections implies an assumption that COTS-driven mortalities are captured by varying background mortality. Incorporating COTS dynamics is a key area of future work to align with GBR management priorities (2, 40). Additionally, macroalgal competition and ocean acidification effects are not explicitly modelled, though these processes may become increasingly important drivers of coral reef dynamics under future climate scenarios (41).

Representation of natural adaptation mechanisms in CoralBlox also has several important limitations. The model represents coral thermal tolerance through truncated normal distributions of DHW tolerances for each functional group and size class, but this statistical approach simplifies the complex genomic underpinnings of heat resistance (42). The heritability coefficient – used to represent the proportion of trait variance that is explained by additive genetic effects (fixed at 0.3; (21)) is drawn from limited empirical studies and may not capture the full complexity of inheritance patterns across diverse coral species across the GBR.

The selection process relies on the univariate Breeders’ equation (43, 44), which assumes that the trait being modelled is causal and entirely responsible for the organism’s biological fitness (45). The approach does not fully account for non-linear genomic interactions, epistasis, physico-chemical limits to adaptability or potential trade-offs between thermal tolerance and other fitness components. The model also assumes a stable genetic variance through fixed standard deviations of tolerance distributions, which may not reflect how genetic variance changes under strong selection pressures over time. Furthermore, the connectivity-based weighting system for inheritance represents a simplification of complex larval dispersal and settlement dynamics.

Standing genetic variation and potential de novo mutations, represented in the model as the upper bound relative to each population’s initial mean, allow populations to evolve to higher thresholds of heat stress tolerance, up to a limit of +24 DHW from the initial mean (35). Despite early concerns around potentially biasing model results towards adaptation levels that are implausibly high, it is found that the represented populations do not adapt to that degree prior to population collapse. The approach does, however, lack strong empirical validation and could under or over-estimate a population’s capacity to adapt to climate change. While these simplifications are necessary for computational efficiency, they represent areas where future work could enhance biological realism as our ability to accurately parameterise and predict coral adaptation improves.

The calibration process demonstrates that CoralBlox can produce credible ecosystem behavior and trajectories. It is acknowledged, however, that the present work is limited to a single “best fit”, contingent on the initial state. The calibration itself is limited by incomplete representation of disturbances such as cyclone damage, which is currently handled via forcing data. Ensemble approaches will be adopted in the future to assist in identifying relevant and critical areas of parameter space (46, 47) and facilitate a Value-of-Information assessment. A multi-model ensemble approach could also be adopted in future to account for structural uncertainties and gain increased confidence in projections (48–50).

### 3.2 Representing intervention activities

Seeding of thermally enhanced corals can be represented by adjusting the DHW tolerance distributions via the simple distribution average approach, weighted by their contribution to area covered. The mixing of the thermally enhanced population and the inheritance of those enhancements are captured by the approach outlined in the “Natural adaptation, selection and inheritance” section in Materials and methods. Efforts to reduce DHW influences locally or regionally (e.g., through geoengineering) can also be represented by adjusting DHW values.

Future work will focus on incorporating these missing disturbances, refining the representation of adaptation mechanisms, and expanding the model’s capacity to represent intervention activities. The ability of CoralBlox to reproduce past ecological trends demonstrates its potential to serve its purpose as a rapid assessment tool for exploration of management options.

## 4. Materials and Methods

Various approaches to modelling coral dynamics exist in literature, each serving different practical purposes. A common approach is to use Leslie or transition matrices, representing transitions between different species’ life stages and mortality through matrix operations (51–53). ODE approaches are common in ecological modelling, capturing continuous dynamics by explicitly modelling instantaneous rates of growth, mortality, and species interactions (54, 55). However, without augmentation, ODEs can fail to represent ecological and environmental stochasticity and uncertainty. Various alternative formulations seek to address this, including nonlinear stochastic models (56, 57). Here, we present a computationally efficient approach to modelling coral population dynamics that balances complexity, uncertainty representation, and computational time.

CoralBlox represents a reef’s coral cover through a set of discretized distributions of coral sizes for each functional group. Each discretized element of these distributions is updated at each timestep to account for distinct ecological and environmental processes (see Figure 1B). Here, a timestep represents a one-year cycle. These processes comprise coral growth and mortality dynamics, coral reproduction, natural adaptation to heat stress, and acute disturbance events that impact coral population trajectory (see Figure 1A). The disturbances include heat-stress-related mortality and cyclone-related mortality. The order of events comes from the assumption that coral spawning occurs between October and November (with the full moon), the peak cyclone season is between January and March, bleaching occurs between February and March, and mortality events occur between April and August.

The model represents reef systems in a spatially lumped manner and agnostic to spatial scale. Each location of concern is represented by a polygon that either represents an entire reef or a discretized area of a reef. In GBR-scale simulations, reefs are represented individually. Finer-scale simulations can be conducted with greater spatial detail, for example, by discretizing reef areas based on their geomorphic and benthic properties to provide a more explicit representation of the reef seascape (58). In the rest of this section, we favor the term “location” over “reef” given this spatial agnosticism.

Overcropping by macroalgae is currently not represented, nor are the potential negative effects of ocean acidification. These disturbances are nonetheless important drivers of coral population dynamics and are planned to be implemented when suitable representations are established.

### 4.1 Model structure and parameterization

CoralBlox represents five distinct coral functional groups, namely tabular *Acropora* (TA), corymbose *Acropora* (CA), corymbose non-*Acropora & Pocillopora* (CNA), small massives representing a mix of encrusting and submissive corals (SM), and large massives such as large porites (LM) indexed by *g ∈* {1, …, *G*} ⊂ ℤ^+^, *G* = 5, in that order. Each coral colony is approximated by a circle and the entire population for each functional group is represented by a discretized diameter distribution. The minimum and maximum diameter for each functional group *g* are 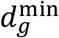 and 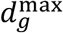, respectively. The range of valid diameters for each functional group is divided into seven non-equal size classes, labeled *z ∈* {1, …, *Z*} ⊂ ℤ^+^, *Z* = 7. Each size class *z* for functional group *g* has lower and upper bounds 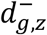 and 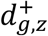, such that if a coral colony that belongs to functional group *g* and has diameter 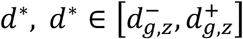, it belongs to that functional group’s size class *z*. This division into size classes enables us to account for size-dependent ecological dynamics, such as mortality and growth.

For each one-year timestep (*t*), the coral cover for each location *l, l ∈* {1, …, *L*} ⊆ ℤ^+^, is independently updated. Locations are assumed to have a habitable area *H*_*l*_, corresponding to the proportion of their total area where corals can live. The coral cover relative to the habitable area *H*_*l*_ for functional group *g* and size class *z* is denoted by *C*_*t*;*g,z,l*_ *∈* [0,1] ⊂ ℝ^+^. The coral cover for a location *l*, summed up across all functional groups and size classes, is denoted by *C*_*t*;*l*_ *∈* [0,1] ⊂ ℝ^+^. Values of *C*_*t*;*l*_ = 1 indicate that the entire habitable area is taken up by coral. The available space at a location *l* and timestep *t* is:

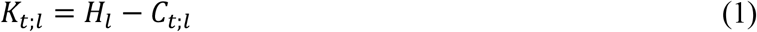

Growth and mortality regional scale factors are included to represent regional dynamics. These are denoted as Ξ_*g,ℓ*_, and ϒ_*g,ℓ*_ for growth and mortality, respectively, and are calibrated with observations across twelve *reef groups, ℓ* = 1, …,12. These reef groups encompass the entire GBR and were built upon bioregions (59, 60) defined by the Great Barrier Reef Marine Park Authority (GBRMPA), clustered together based on spatial proximity to ensure the minimum amount of data for calibration and test sets. Refer to the “Model calibration and testing” section in Materials and Methods for more details.

The coral cover for each location is computed at the functional group and size class level and then summed. Coral cover *C*_*t,l*_ for the location *l* at the end of timestep *t* is given by:

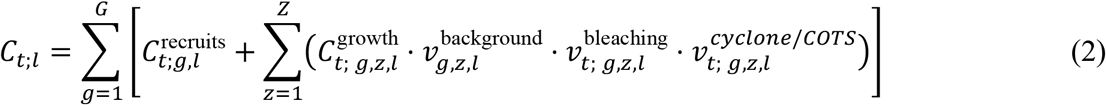

where the recruits cover 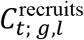 represents the increase in coral cover resulting from reproduction (see “Coral reproduction” section in Materials and Methods) and the growth cover 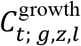 accounts for the coral cover after a growth event (see the “Coral growth” section in Materials and Methods). The survival rates *v* can be written in terms of their respective mortality rates as *v* = 1 − *m*. The *background* mortality rate, 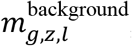, account for the mortality under stable conditions adjusted by the regional scale factor:

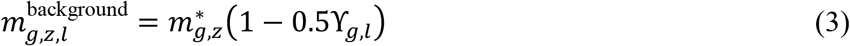

where 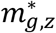 is the base background mortality value. The bleaching mortality rates, 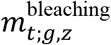, are derived from the observed relationships between heat stress and coral mortality (see section “Coral bleaching and mortality”). Although CoralBlox does not explicitly model cyclone or COTS related damage, the cyclone/COTS survival rates 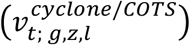 were derived from LTMP historical data for the purpose of assessing how well coral dynamics are captured.

### 4.2 Coral growth

As mentioned, CoralBlox represents each functional group as discretized distributions of coral diameter (Figure 1B). These distributions are composed of *blocks* and indexed by *b* = 1,2, …. Each of these blocks represents the number of corals, *λ*_*g,z,b*_, whose diameter lies within the interval 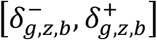. The indices *g, z, b* indicate that this is the *b*-th *block* from functional group *g* and size class *z*. We say that a *block b* belongs to a size class *z* if 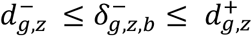 and 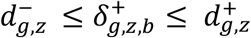. A *block* always belongs to one, and only one, size class.

To calculate the coral cover from this distribution of diameters, consider the area of a coral colony, approximated by the area of a circle with diameter *x*:

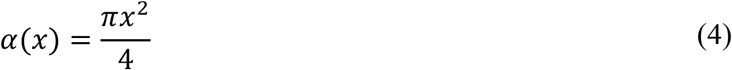

Given a diameter density function *λ*(*x*), representing the number of corals per unit of size with diameter *x*, the area *C*(*x*_*i*_, *x*_*f*_) covered by corals with diameter between *x*_*i*_ and *x*_*f*_ is given by:

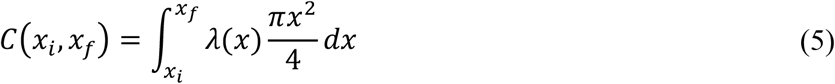

Since the diameter density of each coral block is constant, we can use Eq. (5) to calculate the area covered by the corals within each block:

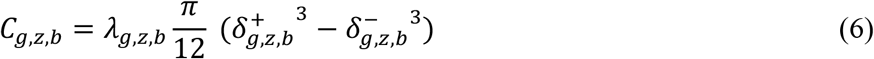

Each coral colony has a linear extension, *γ*_*g,z*_, that represents change in diameter for each coral colony over a year, further scaled at each location by the regional growth factor Ξ_*g,l*_. Linear extensions assume constant distinct values for corals within the same functional group *g* and size class *z*. Since the coral cover for each location is spatially constrained by that location’s habitable area, when the coral cover is high, competition for space takes place. Conversely, when coral cover is low, growth is unconstrained and faster. To address both effects, at each timestep we make use of spatial competition coefficients (*κ*_*t*;*l*_), based on the location’s available space *K*_*t*;*l*_ and habitable area *H*_*l*_ and on the regionally-scaled linear extensions *γ*_*g,z*_Ξ_*g,l*_, and growth acceleration coefficients (*ζ*_*t*;*l*_), based on the location’s available space *K*_*t*;*l*_. Refer to section “Coral growth and mortality” in the Supplementary Materials for more details on how these growth coefficients are calculated, including why *κ*_*t*;*l*_ must be solved from the regionally-scaled linear extensions rather than *γ*_*g,z*_ alone. Rather than switching abruptly between the two coefficients, growth transitions smoothly with cover: locations with low relative cover (*C*_*t*;*l*_ < 0.05) grow solely according to *ζ*_*t*;*l*_, locations with high relative cover (*C*_*t*;*l*_ ≥ 0.80) grow solely according to *κ*_*t*;*l*_, and locations with intermediate cover use a linear blend of the two. The actual growth rate used at each timestep is given by:

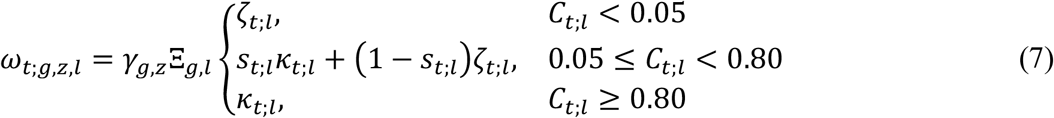

where *s*_*t*;*l*_ = (*C*_*t*;*l*_ − 0.05)/0.75 is the transition weight between the two regimes.

A growth event is represented as the displacement of each coral block in the diameter space, where the growth rates act as translational velocities. The growth rates are always smaller than their correspondent size class widths, 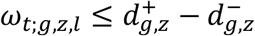, for all *g, z, l* and *t*. Hence after a growth event, we have one of the three following situations for each *block*: (i) the entire *block* remains within size class *z*; (ii) the entire *block* has moved to size class *z* + 1; (iii) part of the *block* is within size class *z* and part is within size class *z* + 1. In the latter case, once the growth event ends, the *block* is split into two, each belonging to only a single size class and having a distinct constant diameter density, where the diameter density of the new *block* that lies within size class *z* + 1 is calculated so that to conserve the number of coral colonies.

Settlers resulting from the reproduction process join the coral population in the following timestep as a block of corals in the smallest size class (*z* = 1). Hence, at each growth event we assume that all corals from the first size (*z* = 1) class move to the next size class (*z* = 2). We also assume that corals in the last size class (*z* = 7) have a constant growth rate of 0.

After a growth event, the total cover 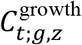, corresponding to corals belonging to the functional group *g* and size class *z*, can be found by using Eq. (6) summed over all its *block****s***:

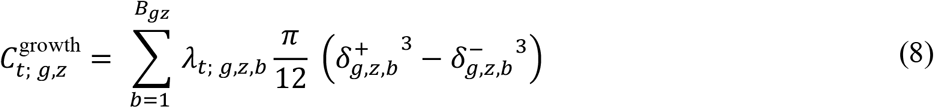

*B*_*g,z*_ is the number of *blocks* within size class *z* for functional group *g* and 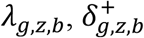 and 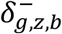 are, respectively, the diameter densities, upper bounds and lower bounds for each coral block after the growth event has happened.

### 4.3 Coral reproduction

Fecundity is first considered within the scope of different coral functional groups and size classes, without consideration of environmental factors. We define *ϕ*_*g,z*_ as the number of larvae produced per colony, which is then converted into the number of larvae per m^2^ for each size class and functional group. The number of larvae per colony is calculated as a function of colony area based on models derived from observational data (61):

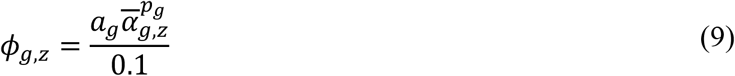

Where *a*_*g*_ and *p*_*g*_ are fecundity parameters from (61) and 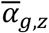 is the mean colony area for the functional group *g* and size class *z*. The number of larvae produced by each functional group and location, *ϕ*_*g,l*_, is then determined by multiplying the location coral area and the summation of all size classes:

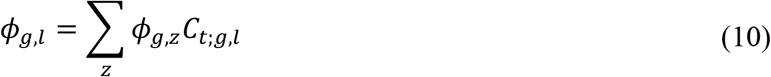

Spawning and recruitment processes represent the release of gametes by broadcast spawning corals into surrounding waters where they fertilize and form larvae, and the proportion of larvae that survive, settle and are recruited as juvenile coral polyps. Larvae may be carried off by currents and never settle, settle on the same site at which they were spawned, or travel to nearby sites depending on ocean currents and other factors, including larval development, mortality, competing organisms, habitat suitability and environmental factors (58). In this study, coral larval dispersal and settlement are represented using connectivity matrices derived from a connectivity model. This model comprises a particle tracking simulator, OceanParcels (62), with velocity outputs from a high-resolution hydrodynamic model, RECOM (63), an implementation of SHOC (64) as input. RECOM velocities are nested within a 1 km resolution hydrodynamic model of the GBR (version 2.0) of the eReefs hydrodynamic model (46), also an implementation of SHOC (47). Nesting RECOM within a lower-resolution model allows particles (virtual larvae) that travel outside RECOM boundaries and potentially re-enter RECOM domain to still be tracked. Details of the creation of the connectivity datasets can be found in (65).

Using these methods, connectivity matrices are generated for several days and years of spawning to reflect the innate stochasticity of the spawning process. The elements *P*_*l*_′_,*l*_ of a connectivity matrix represent the likelihood of larvae moving from location *l*^′^ to *l*, where *l*^′^, *l* = 1, …, *L*. Note that, as larvae can also be carried outside of the domain or die, the rows of *P* need not sum to 1. The number of potential settlers (or larval pool) *R* at each location is then calculated by multiplying the connectivity matrix (averaged over the modelled days and years to capture variation) by the fecundity scope,

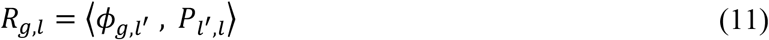

The number of settlers recruited per functional group and location is then modelled following (39, 66), which treats recruitment as a stochastic process based on a Poisson distribution. A rate parameter Λ_*g,l*_ is calculated as the settler density in settlers/m^2^, where Λ_*g,l*_ itself is estimated using a Beverton-Holt function, following (39):

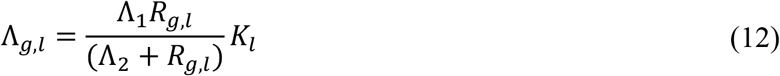

where Λ_1_ is the maximum achievable density, Λ_2_ is the larval stock required to produce 50% of the maximum settlement, and *K*_*l*_ is the space available for coral cover at the location *l*.

### 4.4 Coral bleaching and mortality

Susceptibility to coral bleaching is assessed independently for each location. The relationship between heat stress induced coral bleaching and mortality is based on maximum annual DHW values obtained from the National Oceanic and Atmospheric Administration (NOAA) and observations of coral mortality on the Great Barrier Reef (67).

The coral population is conceptualized as having a truncated normal distribution of heat stress tolerance *F*_*t*; *g,z,l*_ for each functional group (*g*), size class (*z*) and location (*l*). This distribution indicates the proportion of the population that is susceptible to bleaching mortality at a given DHW value. Denoting a truncated normal distribution *TN*(*μ, σ, f*^−^, *f*^+^), where *μ* and *σ* are the mean and standard deviations of the underlying normal distribution, and *f*^−^ and *f*^+^ are the lower and upper truncation bounds, respectively, we can write *F*_*t*; *g,z,l*_ as:

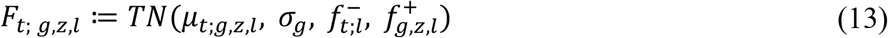

The upper bound 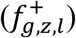 is a hard ceiling fixed at 24 DHW-weeks above each population’s initial mean tolerance (*μ*_*t*=0;*g,z,l*_). The lower bound 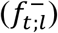 is not a fixed value but carries a one-timestep memory of each location’s own bleaching history:

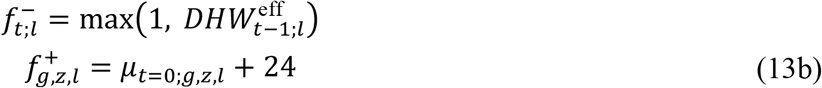

where 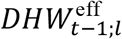 is the effective, depth-attenuated DHW at location *l* during the previous timestep (see Eq. (14) below). Taking the maximum with 1 means 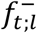 reduces to a floor of 1 DHW-week whenever the previous timestep’s effective DHW was itself below that floor, representing a lack of bleaching susceptibility below that level; following a timestep with effective DHW above 1, the threshold instead tracks how severe the preceding year’s heat stress was, so a location that has already been through a heat-stress event needs at least as much heat stress the following year to affect a further proportion of the population. The distribution shape is informed by data from (67) and analyses on the heritability of coral traits (65). For each functional group, the standard deviations (*σ*_*g*_) are assumed to be identical for all size classes and locations, and are fixed throughout the entire model run to represent stable genetic variance (65). Table S5 shows the values used for all *σ*_*g*_. The mean values (*μ*_*t*;*g,z,l*_) are initially the same for all locations but vary between functional group and then updated at each timestep to account for natural adaptation. Table S4 shows the initial values adopted for all *μ*_*t*=0;*g,z,l*_.

At each timestep, the proportion of the population affected by bleaching for each functional group, size class and location, denoted by *β*_*t*;*g,z,l*_, is estimated based on *F*_*t*;*g,z,l*_, evaluated at an effective, depth-attenuated DHW value rather than the raw surface DHW. This effective DHW 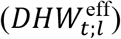 is obtained by attenuating the surface DHW value observed at location *l* (*DHW*_*t*;*l*_) with depth (*h*_*l*_), using the Beer-Lambert law of light attenuation as a proxy for the dissipation of heat stress with depth, following the qualitative depth-refuge relationship reported in (68):

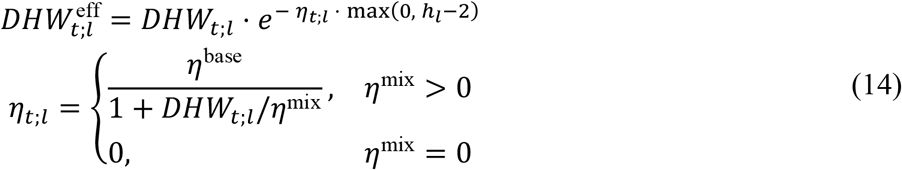

The attenuation coefficient (*η*_*t*;*l*_) is itself a function of a base attenuation coefficient (*η*^base^, in m^−1^) and a mixing factor (*η*^mix^, in DHW), both of which are treated as free parameters and estimated by calibration jointly with the other model parameters (see “Model calibration and testing”). In the low-heat-stress regime (*DHW*_*t*;*l*_ ≈ 0), we have *η*_*t*;*l*_ ≈ *η*^base^; as *DHW*_*t*;*l*_ increases, *η*_*t*;*l*_ decreases, so the depth-refuge effect weakens and saturates at higher heat-stress levels, as shown for the calibrated parameter values in Figure S3. A value of *η*^mix^ = 0 represents the special case in which depth provides no refuge at any DHW. The minimum viable depth for all locations is assumed to be 2 meters below surface, where values lower than 2 offer no protection (68). Bleaching mortality is represented by:

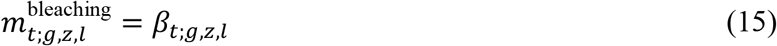

with

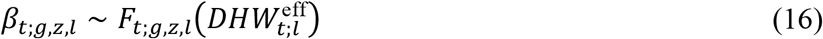

Bleaching mortality is applied to all size classes, as colony size is a weak predictor of bleaching susceptibility (69).

### 4.5 Natural adaptation, selection, and inheritance

After each growth, reproduction and mortality events, the DHW tolerance distributions for each functional group, size class and location need to be updated to capture demographic changes. Here we present each of these updates in the order that they are implemented. Given DHWs vary by location and depth, differences in the distribution of DHW tolerances will emerge over time.

#### 4.5.1 After growth distribution update

Since we assume all corals from size class *z* = 1 move to *z* = 2 after a growth event, we only need to update the DHW tolerance distributions for size classes 2 to 7. The distribution means for each functional group, size class, location and timestep are given by a weighted mean where the weights (*w*) are informed by the proportional coral cover (*C*_*t*;*g,z,l*_) and growth rates (*ω*_*t*;*g,z,l*_):

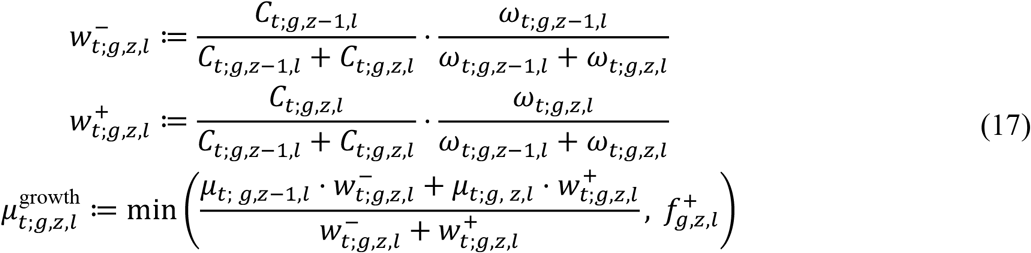

Thus, the updated mean is assumed to be informed by the relative contributions towards the increase in coral cover after growth. Because the ceiling 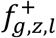 is anchored to each size class’s own initial mean, it can differ between size classes *z* − 1 and *z*; the resulting blend is therefore capped at 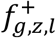, the ceiling of the size class it is now assigned to, so that blending across size classes does not push the mean past that size class’s ceiling. The DHW tolerance distributions after the growth event can be written with Eqs. (13) and (17) as:

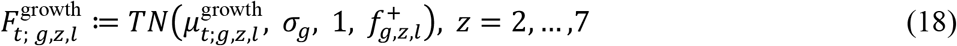

#### 4.5.2 After reproduction distribution update

Heritable adaptation is applied locally at each source location, before larvae disperse: at every location *l*^′^ and for every functional group *g*, the Breeders’ equation (43, 44, 70–72) shifts each reproductively active size class’s tolerance mean from the previous timestep towards its current-timestep value, scaled by the heritability coefficient *h*^2^, and these shifted means are combined across the reproductively active size classes, weighted by their relative cover, into a single tolerance mean representing that location’s contribution to the larval pool:

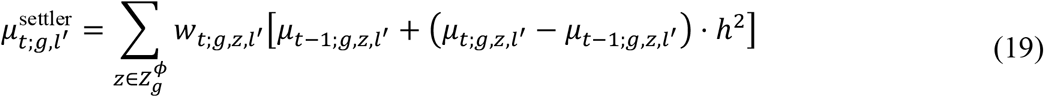

where 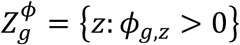 is the set of reproductively active size classes for functional group *g* (see “Coral reproduction” section), and the weights *w*_*t*;*g,z,l*_′ are given by each size class’s relative cover among 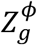:

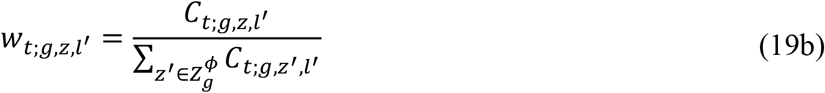

The distribution for the newest generation (*z* = 1) at each sink location *l* is then given by the connectivity-weighted mean of the source contributions 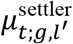, using the same connectivity weights *P*_*l*_′_,*l*_ as Eq. (11):

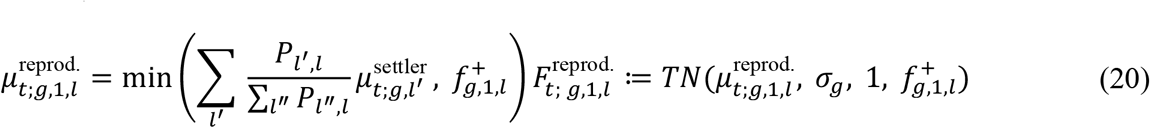

where the heritability coefficient *h*^2^ is assumed to have a nominal value of 0.3. Because the heritable shift in Eq. (19) can push 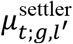 above the source location’s own ceiling, and the connectivity-weighted combination in Eq. (20) can compound this across source locations, 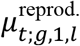 is capped at the sink size class’s ceiling 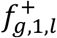 before being used as the mean of the truncated normal.

#### 4.5.3 After bleaching mortality distribution update

After a bleaching mortality event, the new DHW tolerance distribution is simply given by

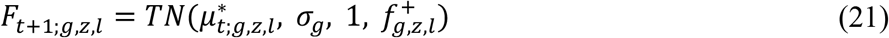

where survivors’ mean is recomputed by truncating below at the current effective DHW (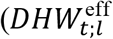, or 1 DHW-week, whichever is greater), since survivors must have had tolerance above the heat stress they were just exposed to:

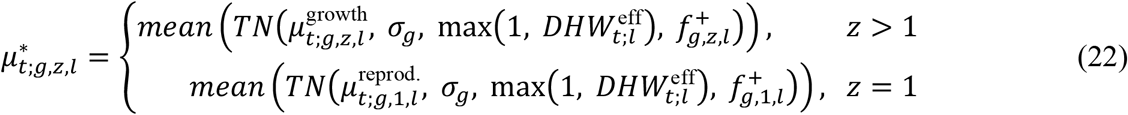

### 4.6 Model calibration and testing

The AIMS Long-Term Monitoring Program has been operating since 1983 and provides observations on the state of different locations within the GBR (6). Observations include total coral cover observed from manta tow surveys and coral composition from fixed site surveys. A subset of the model parameters was selected for calibration to historic observations. Parameters were manually selected for calibration for their influence in growth and mortality dynamics. Some of the calibrated parameters are constant across all reef groups (linear extensions, background mortality rates, initial mean thermal tolerances, and the DHW depth attenuation parameters), while others are resolved separately for each of the twelve reef groups (rate parameters of coral size exponential distribution, growth acceleration parameters, growth regional scale factors, and base mortality regional scale factors). The model parameters were fit to the calibration set using the Adaptive Differential Evolution algorithm (73), implemented in the BlackBoxOptim.jl Julia package (74), and error statistics were calculated for the test set to assess model performance on unseen data. See “Model calibration and testing” section in Supplementary Materials for further details.

Prior to the calibration, reefs were grouped based on ecological similarity and spatial proximity (Figure S1). This split was done programmatically based on the Great Barrier Reef Marine Park Authority’s bioregions (59, 60). These bioregions were grouped together based on spatial proximity to allow a 75%/25% calibration-test split and comparative assessment within each reef group. Refer to the “Data split and reef groups” section of the Supplementary Materials for more details.

**Table 1.**
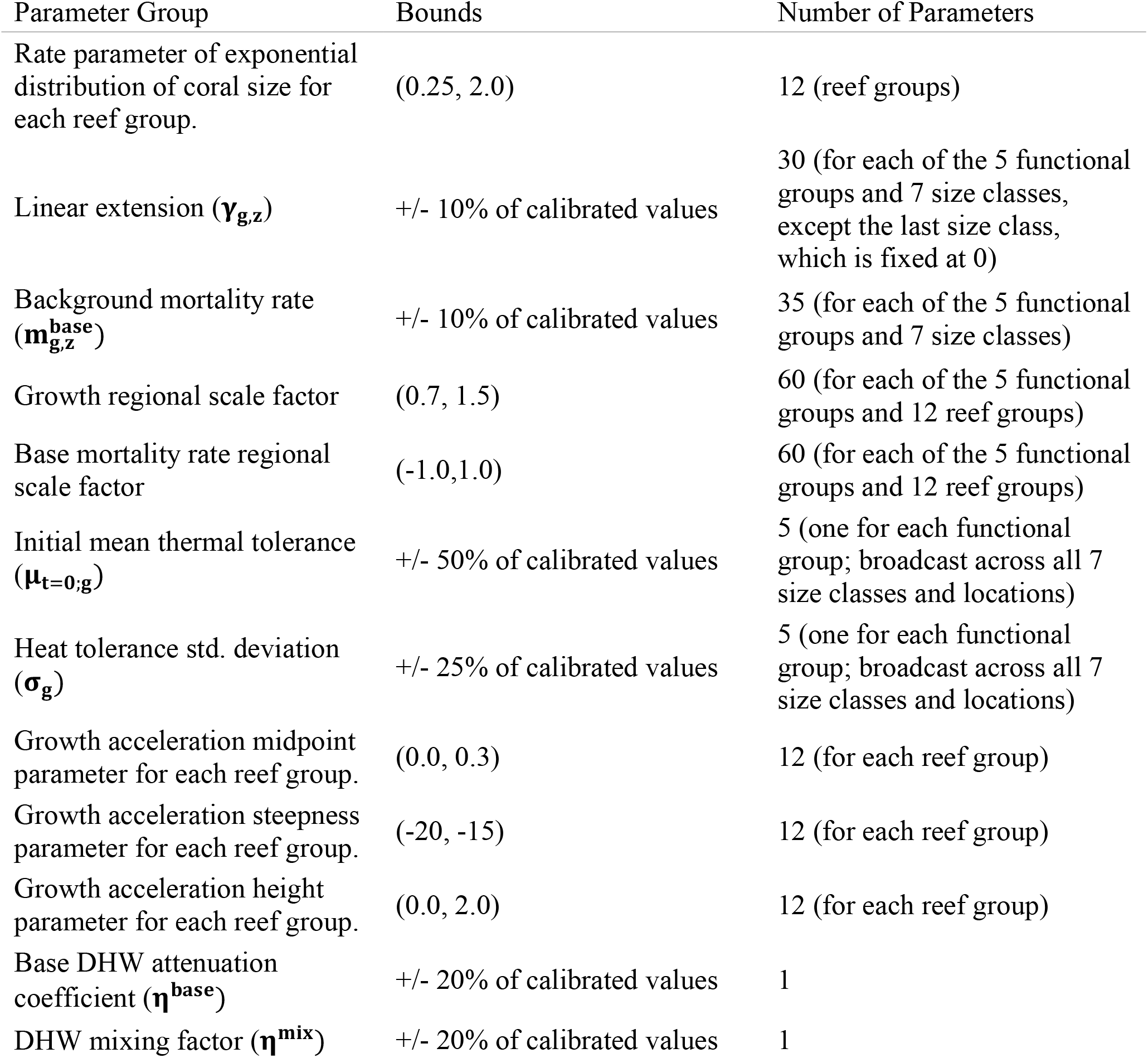
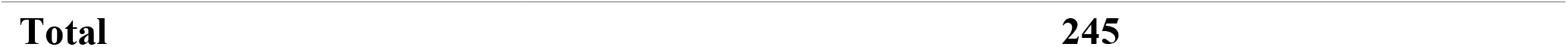
Parameters used in the calibration.

We assess model performance based on modelled and observed relative coral cover for each calibration and test reef separately. For a given reef, let us denote *y* the set of observed relative coral cover values for the year with data, *ŷ* the corresponding modelled relative coral cover values, and 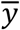 the average across all *y*. The average cover 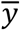 establishes a benchmark providing a minimum but not sufficient performance threshold for CoralBox. The three performance metrics used are i) the difference between modelled and benchmark Root Mean Squared Errors (RMSE) with respect to the observed data, 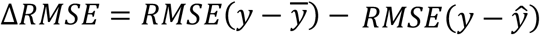, ii) the Spearman Rank Correlation Coefficient (SRCC) between observed and modelled relative coral cover, and iii) the bias, *bias* = *mean*(*ŷ* − *y*). Positive ΔRMSE values indicate reefs where CoralBlox outperforms the benchmark. SRCC values provide an indication of alignment in trends between simulation and observations. Positive bias values indicate CoralBlox overpredicts coral cover on average, negative values indicate underprediction.

Uncertainty in these metrics was quantified via bootstrap resampling, using the Bootstrap.jl Julia package (75). For each location, confidence intervals were obtained by resampling its paired modelled/observed annual values with replacement: locations with five or more years of observations used a circular block bootstrap (block lengths of two to three years, to preserve year-to-year autocorrelation in the error series), while locations with the minimum of four observed years used a standard independent-and-identically-distributed (iid) bootstrap. Both used 2,000 resamples and 95% percentile confidence intervals. The median ΔRMSE, SRCC and bias reported for the calibration and test sets were themselves bootstrapped. This second-stage bootstrap resampled which locations’ point estimates were included and took the median across 2,000 such resamples. Only locations eligible for the block bootstrap were included, so locations with just four years of data contribute a per-location estimate but do not influence the aggregate median’s uncertainty.

We conducted an additional analysis of the model’s performance metrics, focusing on spatial and heat-stress correlations for both dataset splits and both metrics. We categorized spatial correlations into correlations along and across the GBR. For each category, we calculated the SRCC between each reef’s position and each of its performance metrics. For the position along the GBR, we used distance to the northernmost reef, following the definition from (76). For the position across the GBR, we used the haversine distance between each reef and the coast, which was approximated by the minimum of the distances between that reef and each point of the Great Barrier Reef mainland polygon (8). Haversine distances were calculated using the Distances.jl package (77). For the heat-stress correlations, we calculated the SRCC between each reefs’ performance metric and the maximum cumulative thermal exposure, measured in DHW.

AI-assisted tools (Anthropic’s Claude Sonnet 5 and Claude Opus 5) were used during the development of this study to write and refine portions of the code for the CoralBlox and ADRIA model implementations, the calibration pipeline, and the analysis scripts used to compute the performance metrics reported here (ΔRMSE, SRCC, bias, and their bootstrapped confidence intervals). All AI-assisted code was reviewed, tested, and validated by the authors against the model formulation described above prior to use.

## 6. Acknowledgements

We acknowledge the Ngunnawal, Wulgurukaba, Bindal, Whadjuk Noongar, Turrbal and Jagera Peoples of the land on which the authors conducted the model development, analysis and writing for this study. We also acknowledge the Gooreng Gooreng, Gurang, Bailai, Taribelang Bunda, Wunambal Gambera and Gunggandji peoples as the Traditional Owners of the Sea Country where the empirical data for this study was sourced. The work presented here benefitted greatly from discussions and interactions with the EcoRRAP subprogram. We additionally acknowledge the importance of data collected via the AIMS Long Term Monitoring Program in enabling the model evaluation presented here.

The authors would like to thank Yves-Marie Bozec, Arne Adam for advice, comments and suggestions provided during the development of the model and earlier forms of this manuscript. The authors would like to additionally thank Ken Anthony for discussions around coral ecology and early conceptualization of what would eventually become CoralBlox.

## 6.2 Author Contributions

P. Ribeiro de Almeida: Writing, review and editing; data collation and analysis; model and software development.

R. J. Crocker: Writing, model development.

D. Tan: Model and software development, in particular improvements to computational performance; data collation and analysis; writing

K.R. Bairos-Novak: Data analysis and assistance in conceptual development to include representation of natural adaptation, selection, and heritability.

C. J. Ani: Development of the connectivity model. Review.

J. Benthuysen: Review; data collation.

B. Robson: Review

S. Matthews: Review

T. Iwanaga: Writing, review and editing. Model development and changes to incorporate natural adaptation and heritability. Software development for usability and computational performance. Conceptualization of research aims and primary draft. Coordinated manuscript efforts.

## 6.4 Data and Materials Availability

CoralBlox and ADRIA are implemented in the Julia programming language. The source code for CoralBlox is freely and openly available at https://github.com/open-AIMS/CoralBlox.jl and archived at https://doi.org/10.25845/7GZ3-C373 (21). The source code for ADRIA is freely and openly available at https://github.com/open-AIMS/ADRIA.jl and archived at https://doi.org/10.25845/E9VT-4G35 (22). The code used to calibrate the CoralBlox parameters described here is available at https://github.com/open-AIMS/CoralBlox-params-calibration (release v0.5.1; DOI: [to be assigned]) (78). The code to reproduce all analyses and figures in this manuscript is available at https://github.com/open-AIMS/CoralBlox-ADRIA-paper-2026 (DOI: [to be assigned]) (79).

AIMS Long-Term Monitoring Program (LTMP) data, served through the Reef Monitoring dashboard (https://apps.aims.gov.au/reef-monitoring/), was used to calibrate a subset of CoralBlox parameters (6). LTMP manta tow observations were the target for modelled total coral cover outputs and LTMP fixed point photogrammetry community composition data was used as the target for modelled community composition during calibration and testing. These data are licensed under CC-BY 4.0 (https://www.aims.gov.au/cc-attribution); the products used here are based on data supplied by the Australian Institute of Marine Science.

To run the model, we used data from the ReefMod-GBR model (35), freely and openly available at https://github.com/ymbozec/REEFMOD.7.2_GBR. The data as used in this paper is available upon request; please send enquiries for ReefMod Engine to. Further details are available in the CoralBlox calibration repository referenced above.

Mortality rates accounting for mortalities caused by cyclones and/or COTS that are used in the calibration were derived from the AIMS LTMP disturbance data.

Observed annual maximum degree heating weeks (DHWs) data was derived from daily DHWs provided by the NOAA CoralTempv3.1 product at (https://coralreefwatch.noaa.gov/product/5km/). Annual values are based on 1 July from the year prior to 30 June.

Reef and coastline geometries used for mapping and to derive each reef’s distance offshore are from the Great Barrier Reef Features dataset (8), © Great Barrier Reef Marine Park Authority 2014, licensed under CC-BY 4.0 and obtained via the eAtlas data portal (metadata record ac8e8e4f-fc0e-4a01-9c3d-f27e4a8fac3c).

## 6.5 AI Usage

We acknowledge the use of AI-assisted tools (Claude Sonnet 5 and Claude Opus 5, Anthropic) in preparing this manuscript: drafting and revising portions of the text, generating the Quarto code used to render the manuscript and its figures, and producing the plots and performance metrics presented in the Results and Supplementary Materials. All AI-assisted content was reviewed and verified by the authors.

## 7. Supplementary Materials

### 7.1 Parameter values

**Heritability = 0.3**

### 7.2 Tables

**Table S1.**
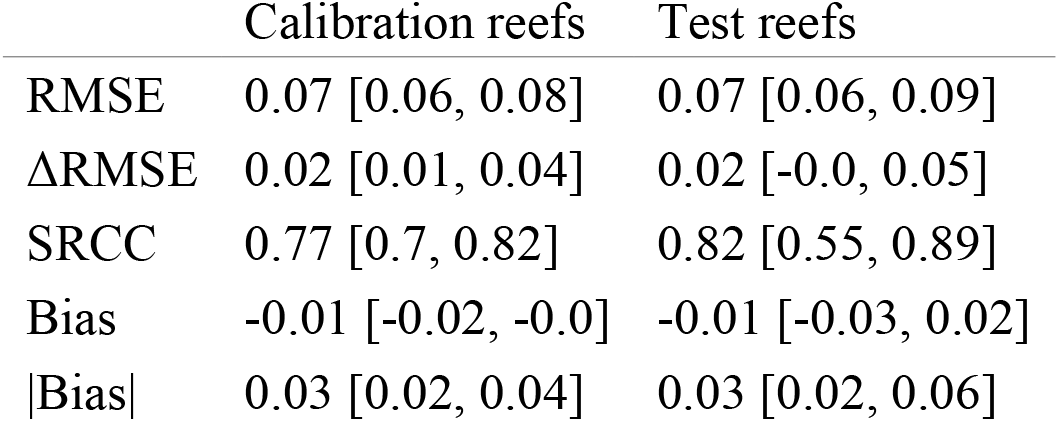
Bootstrapped median performance metrics and 95% confidence intervals for calibration and test reefs.

**Table S2.** Bootstrapped SRCC with 95% confidence intervals between performance metrics and positions along and across the GBR, as well as maximum DHW.

|  | Along the GBR | Across the GBR | Maximum DHW |
| --- | --- | --- | --- |
| ΔRMSE Calibration | 0.38 [0.19, 0.6] | -0.4 [-0.64, -0.22] | -0.42 [-0.63, -0.23] |
| ΔRMSE Test | 0.44 [0.2, 0.79] | -0.38 [-0.75, -0.13] | -0.53 [-0.9, -0.28] |
| SRCC Calibration | 0.24 [0.02, 0.49] | -0.26 [-0.5, -0.06] | -0.48 [-0.67, -0.3] |
| SRCC Test | 0.4 [0.15, 0.78] | -0.34 [-0.71, -0.05] | -0.48 [-0.81, -0.25] |

**Table S3.** Size class bin edges (d^−^ to d^+^) for each functional group (columns) and size class (rows) in cm.

| $z$ | TA ($g = 1$) | CA ($g = 2$) | NAC ($g = 3$) | SM ($g = 4$) | LM ($g = 5$) |
| --- | --- | --- | --- | --- | --- |
| 1 | 2.5 to 7.5 | 2.5 to 7.5 | 2.5 to 7.5 | 2.5 to 5.0 | 2.5 to 5.0 |
| 2 | 7.5 to 12.5 | 7.5 to 12.5 | 7.5 to 12.5 | 5.0 to 7.5 | 5.0 to 7.5 |
| 3 | 12.5 to 25.0 | 12.5 to 20.0 | 12.5 to 20.0 | 7.5 to 10.0 | 7.5 to 10.0 |
| 4 | 25.0 to 50.0 | 20.0 to 30.0 | 20.0 to 30.0 | 10.0 to 20.0 | 10.0 to 20.0 |
| 5 | 50.0 to 80.0 | 30.0 to 60.0 | 30.0 to 40.0 | 20.0 to 40.0 | 20.0 to 40.0 |
| 6 | 80.0 to 120.0 | 60.0 to 100.0 | 40.0 to 50.0 | 40.0 to 50.0 | 40.0 to 50.0 |
| 7 | 120.0 to 160.0 | 100.0 to 150.0 | 50.0 to 60.0 | 50.0 to 100.0 | 50.0 to 100.0 |

**Table S4.** Calibrated values for initial means (μ_t=0,g_) of the truncated normal distributions (TN) of heat tolerance for each functional group, in Degree Heating Weeks (DHW). As with σ_g_ (Table S5), μ_t=0_ is calibrated per functional group and broadcast across all seven size classes and locations, not calibrated independently per size class.

| Functional Group | Initial Mean ($\mu_{t=0,g}$) |
| --- | --- |
| TA ($g = 1$) | 5.63 |
| CA ($g = 2$) | 6.12 |
| NAC ($g = 3$) | 6.73 |
| SM ($g = 4$) | 3.78 |
| LM ($g = 5$) | 3.58 |

**Table S5.** Standard deviation (σ_g_) of the truncated normal distributions of heat tolerance (in DHW) assumed for each functional group.

| Functional Group | Standard Deviation ($\sigma_g$) |
| --- | --- |
| TA ($g = 1$) | 2.50 |
| CA ($g = 2$) | 3.65 |
| NAC ($g = 3$) | 2.61 |
| SM ($g = 4$) | 4.18 |
| LM ($g = 5$) | 4.15 |

**Table S6.** Calibrated base mortality rates 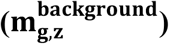 for each functional group (columns) and size class (rows).

| $z$ | TA ($g = 1$) | CA ($g = 2$) | NAC ($g = 3$) | SM ($g = 4$) | LM ($g = 5$) |
| --- | --- | --- | --- | --- | --- |
| 1 | 0.34 | 0.25 | 0.24 | 0.24 | 0.31 |
| 2 | 0.21 | 0.14 | 0.14 | 0.09 | 0.09 |
| 3 | 0.23 | 0.10 | 0.09 | 0.05 | 0.05 |
| 4 | 0.21 | 0.14 | 0.09 | 0.03 | 0.03 |
| 5 | 0.17 | 0.12 | 0.08 | 0.01 | 0.01 |
| 6 | 0.15 | 0.10 | 0.09 | 0.02 | 0.01 |
| 7 | 0.13 | 0.12 | 0.08 | 0.03 | 0.03 |

**Table S7.** Calibrated linear extensions (γ_g,z_) for each functional group (columns) and size class (rows) in cm/year.

| $z$ | TA ($g = 1$) | CA ($g = 2$) | NAC ($g = 3$) | SM ($g = 4$) | LM ($g = 5$) |
| --- | --- | --- | --- | --- | --- |
| 1 | 2.21 | 2.19 | 1.58 | 1.05 | 1.08 |
| 2 | 4.62 | 2.58 | 1.52 | 0.73 | 0.67 |
| 3 | 4.53 | 2.51 | 1.36 | 0.68 | 0.68 |
| 4 | 5.74 | 2.57 | 1.82 | 0.85 | 0.85 |
| 5 | 6.05 | 2.83 | 1.82 | 1.21 | 1.33 |
| 6 | 7.02 | 3.28 | 1.49 | 1.27 | 1.27 |
| 7 | 0.00 | 0.00 | 0.00 | 0.00 | 0.00 |

**Table S8.** Calibrated growth regional scale factors (Ξ_g,ℓ_) for each reef group (rows) and functional group (columns).

| $\ell$ | TA ($g = 1$) | CA ($g = 2$) | NAC ($g = 3$) | SM ($g = 4$) | LM ($g = 5$) |
| --- | --- | --- | --- | --- | --- |
| 1 | 1.03 | 0.72 | 1.40 | 0.70 | 0.70 |
| 2 | 1.08 | 1.10 | 1.27 | 0.78 | 1.44 |
| 3 | 1.08 | 1.24 | 1.11 | 0.70 | 0.77 |
| 4 | 0.98 | 1.37 | 0.74 | 1.38 | 1.50 |
| 5 | 0.72 | 0.90 | 0.73 | 1.24 | 1.18 |
| 6 | 0.88 | 0.70 | 0.89 | 1.09 | 0.79 |
| 7 | 0.84 | 0.88 | 0.86 | 0.87 | 0.70 |
| 8 | 0.90 | 0.85 | 0.75 | 0.78 | 1.09 |
| 9 | 1.06 | 1.00 | 0.87 | 1.03 | 0.76 |
| 10 | 1.08 | 0.98 | 1.18 | 1.37 | 0.83 |
| 11 | 0.71 | 0.79 | 0.78 | 0.74 | 0.72 |
| 12 | 1.06 | 0.86 | 0.82 | 0.70 | 0.91 |

**Table S9.**
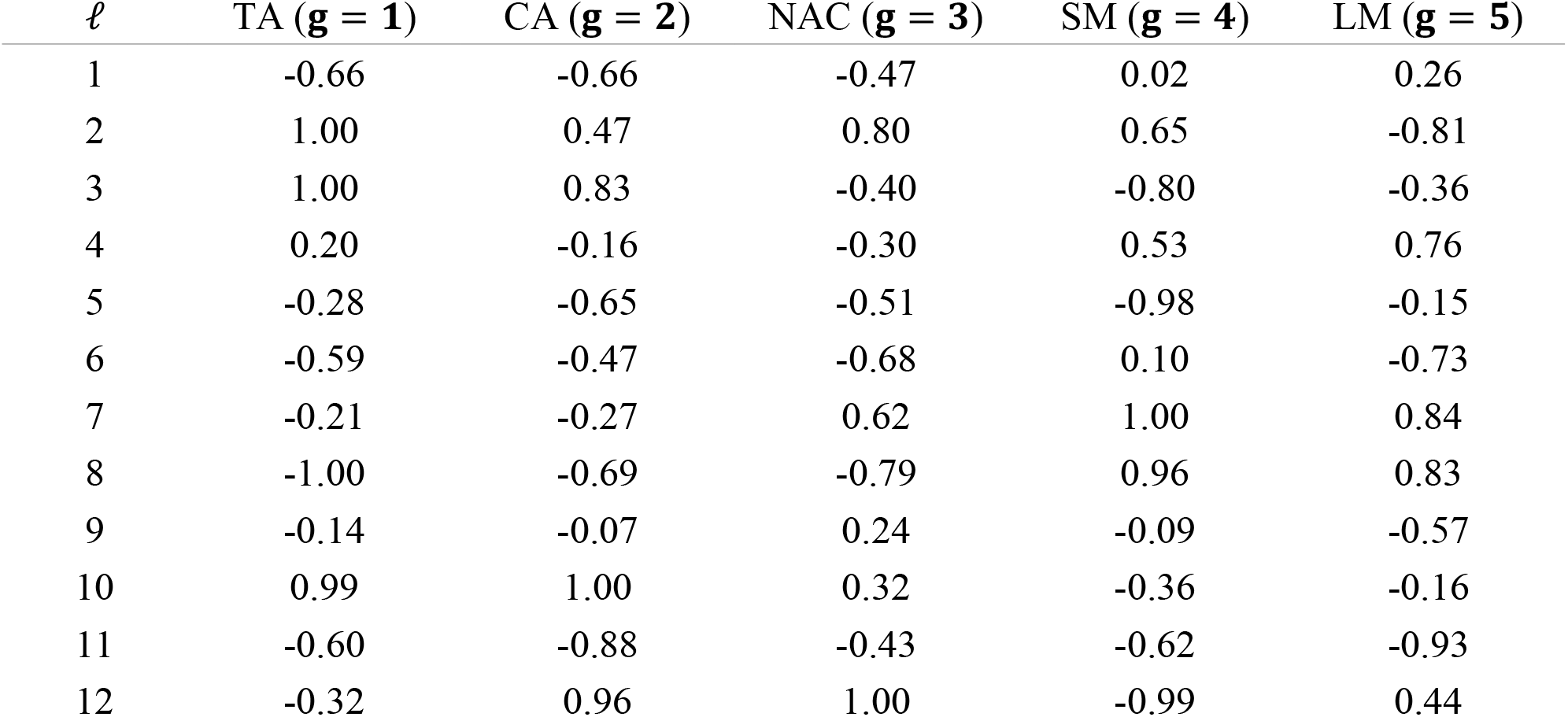
Calibrated background mortality scale factors (ϒ_g,ℓ_) for each reef group (rows) and functional group (columns).

**Table S10.**
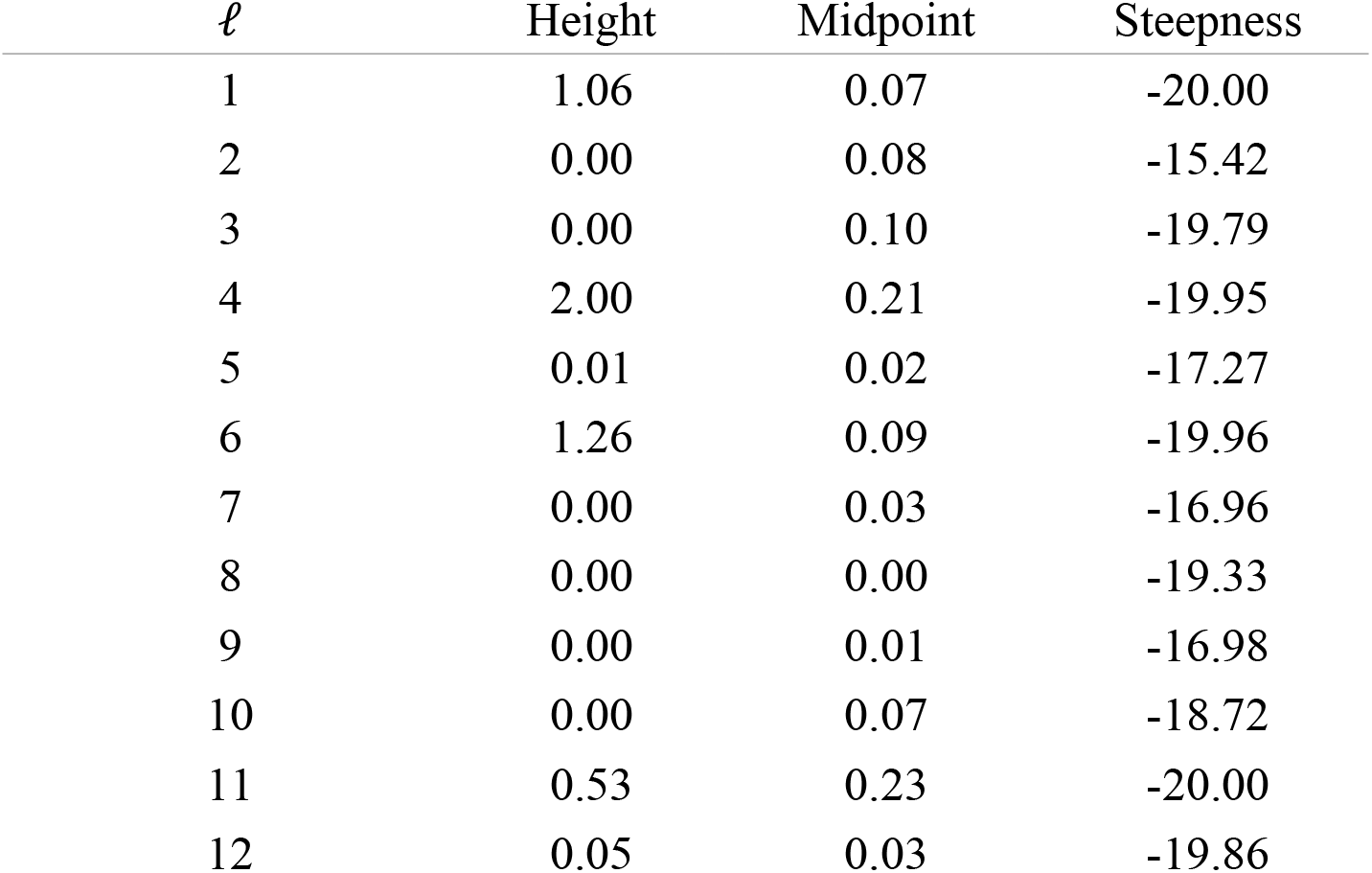
Calibrated growth acceleration parameters (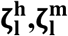 and 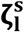) for each reef group (rows).

### 7.3 Coral block movement

As detailed in the Materials and Methods section, CoralBlox represents corals by chunks (*blocks*) of a discretized diameter distribution. Each *block b* from functional group *g* and size class *z* is characterized by its lower and upper bounds (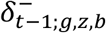 and 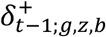) and height (*λ*_*t*−1;*g,z,b*_). A growth event is represented by the movement of each block in the diameter space with speed *ω*_*g,z*_, measured in diameter units per year. When a block moves, we can have one of the following situations:

1. if 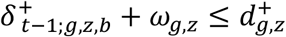, then the block moves only within the current size class.
2. If 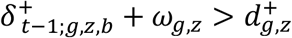 and 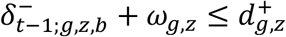, then the block’s upper bound moves from one size class to the next one but the lower bound doesn’t.
3. If 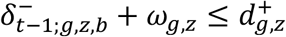 then the entire block moves from one size class to the next one.

Where 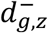 and 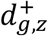 are the lower and upper bounds of size class *z* from functional group *g*. When a *block* moves within the same size class, the lower and upper bounds after a growth event are easily determined. On the other hand, when a block crosses the boundary between two consecutive size classes, we need to be careful since two distinct growth rates are acting at the same time on the same block. This situation is similar to a stream of water flowing between two pipelines of different diameters. The resulting water flow speed changes based on the pipeline diameter, but the amount of water as input in the system is constant. In the same way, when a coral density *block* moves between size classes, its speed and height (diameter density) change but the number of coral colonies represented by that block remains constant.

This section analyzes the movement of a coral density *block* in detail. We assume that the movement begins at timestep *t* − 1 with the block entirely within size class *z*. The time it takes for the block to move is Δ*t* = 1, measured in years. A coral block can only move forward within its current size class or grow into the next larger size class. This represents a unidirectional progression through size classes that accounts for coral growth and does not model size reduction, which would require bidirectional movement between size classes.

In case 1, the bounds for the *block* after the growth event, at timestep *t*, are given by:

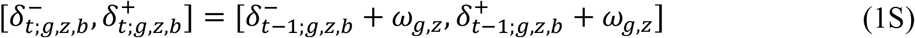

Note that, since Δ*t* = 1, we have *ω*_*g,z*_Δ*t* = *ω*_*g,z*_ on the left-hand side of Eq. (1S), so there is no dimension mismatch. In cases 2 and 3 we need to analyze the lower and upper bounds movements separately and then adjust the coral block density accordingly to conserve the number of corals. In both cases, the upper bound moves with speed *ω*_*g,z*_ until it reaches the current size class upper bound 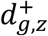, then it starts to move with speed *ω*. The total displacement of the upper bound Δ*δ*^+^ is given by:

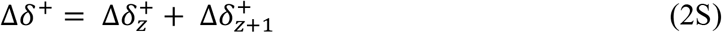

where 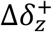 and 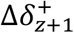 correspond to the movement within size classes *z* and *z* + 1, respectively. The indices *g, b* are omitted for simplicity. At the end of a timestep, the upper bound is given by:

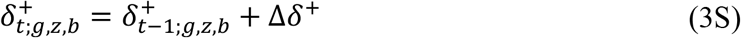

Our goal is to determine Δ*δ*^+^. Let us denote Δ*t*_*z*_ the time it takes for the *block*’s upper bound to reach 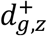 and Δ*t* the time it takes to go from there to 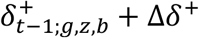, so that

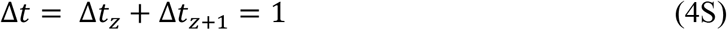

The first term of Eq. (2S) is known, since one of the assumptions is that the coral block’s upper bound crosses the upper bound of size class *z*:

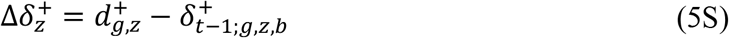

Hence, using Eq. (5S) and the fact that for a uniform linear movement speed equals spatial displacement divided by time interval, we have:

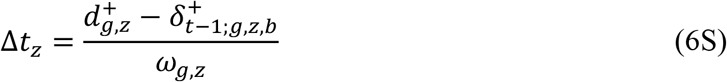

Using Eq. (4S) we have Δ*t*_*z*+1_ = 1 − Δ*t* and, since 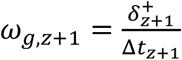, we have

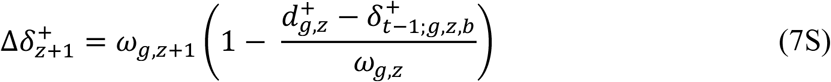

And putting Eqs. (2S), (5S) and (7S) together, the total displacement of the upper bound is given by:

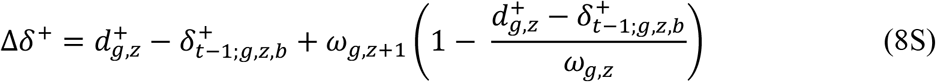

In case 2, where only the upper bound crosses the interface between size classes, at the end of the timestep we split that coral block in two, the first with bounds 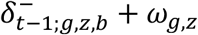 and 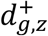 and the second with bounds 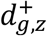 and 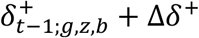.

The density of the first block remains the same as before, *λ*_*g,z,b*_. With the first block’s width and density, we can determine the number of coral colonies that moved to the next size class by:

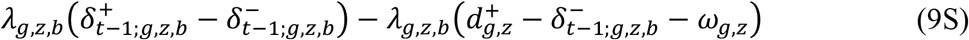

and we can use that to find the density of the coral block that moved to the next size class:

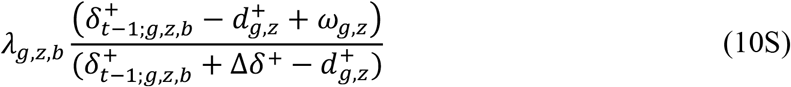

In case 3, where both lower and upper bounds move to the next size class, the coral block’s bounds are given, at the end of the timestep, by 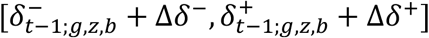 where Δ*δ*^−^ is, in analogy with Eq. (8S), by:

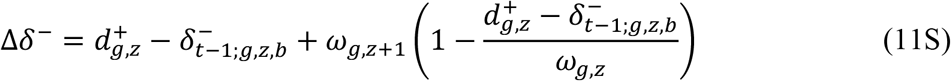

and in this case the new coral density is given by:

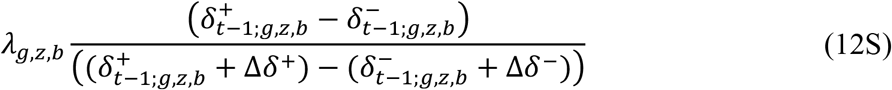

### 7.4 Coral growth and mortality

The growth rates used at each timestep are based on the regionally-scaled linear extensions *γ*_*g,z*_Ξ_*g,l*_ and are rescaled by the coral growth and the spatial competition coefficients, as per Eq. (7). This section presents details on these two coefficients calculations.

#### 7.4.1 Low cover growth acceleration

When coral cover at a location is low, corals are unencumbered by competition and will grow at a faster rate. To account for this, growth acceleration coefficients *ζ*_*l*_, following a sigmoid function, are used to replicate faster growth rates at low cover levels and gradually transition to normal growth rates as cover approaches moderate levels. The *ζ*_*l*_ for each location *l* are given by:

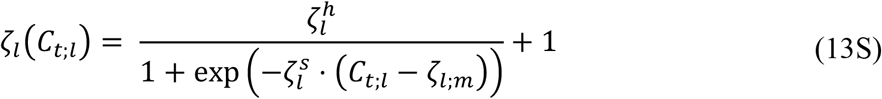

where 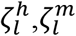 and 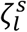 are the coefficient’s height, midpoint and steepness parameters, respectively, and *C*_*t*;*l*_ is the relative coral cover summed across all functional groups and size classes of location *l* at timestep *t*.

#### 7.4.2 Coral growth and spatial competition

The spatial competition coefficients *κ*_*t*;*l*_ are calculated at every timestep as it ensures that the total coral cover never outgrows the habitable area. Although in the growth event at each timestep we move one coral block at a time, *κ*_*t*;*l*_ captures all functional group’s simultaneous growth. Two relevant properties of *κ*_*t*;*l*_ are that 1) it makes the growth rate decrease as the coral cover *C*_*t*−1_ grows and the available space (Eq. (1)) decrease and, 2) when there is no available space left, it makes the growth rate become zero.

To calculate *κ*_*t*_, we follow three steps. First, we find an approximate projected cover 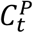 using *γ*_*gz*_Ξ_*g,l*_ as the growth rate, as if there was no spatial constraint. Second, we apply a normalization function to transform 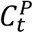 into an adjusted cover 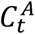 whose values lie between the interval [*C*_*t*−1_, *H*_*l*_]. Third, we find a constant scale factor *κ*_*t*_ such that, when we repeat the same procedure used at the first step to calculate the approximate projected cover, but using *ω*_*gz*_ = *γ*_*gz*_Ξ_*g,l*_*κ*_*t*_ as the growth rate, the resulting scaled cover 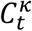 is equal to 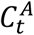.

Note that *γ*_*gz*_ is scaled by Ξ_*g,l*_ *before* this three-step derivation, rather than being applied afterwards to the resulting *ω*_*gz*_. This ordering is necessary for *κ*_*t*_ to retain its defining property: since *κ*_*t*_ is solved so that the growth rate actually applied never lets 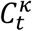 exceed the habitable area *H*_*l*_, that guarantee only holds if the growth rate used to derive *κ*_*t*_ (Eqs. (15S)-(20S)) is the same growth rate the population ultimately grows by. Scaling *γ*_*gz*_ by Ξ_*g,l*_ only after solving for *κ*_*t*_ would decouple the two, allowing 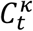 to exceed *H*_*l*_ whenever Ξ_*g,l*_ ≠ 1. The linear extensions *γ*_*g,z*_ appearing in Eqs. (15S) and (18S)-(20S) below are therefore understood to already include this regional scaling, i.e. *γ*_*g,z*_Ξ_*g,l*_.

#### 7.4.3 Projected Cover

To calculate the approximate projected cover 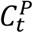 at timestep *t*, we approximate the cover correspondent to all coral blocks inside each size class by an average block, with diameter density given by:

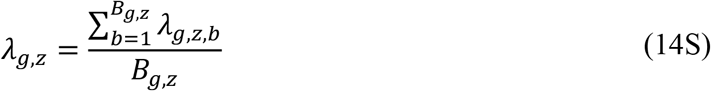

where *B*_*g,z*_ is the number of coral blocks within size class *z* and functional group *g*. We also assume that each coral block will follow a uniform movement. This way, using Eq. (8) we can calculate 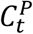 as:

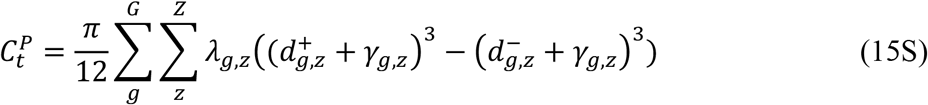

#### 7.4.4 Adjusted Projected Cover

The adjusted projected cover is given by:

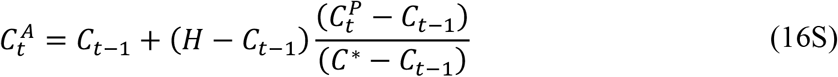

where the maximum projected cover *C*^∗^ is an upper bound to 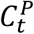. Since there are infinite possible upper bounds for 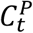, we chose one that balances accuracy and computation speed: *C*^∗^ is obtained by concentrating the entire habitable area *H* into whichever functional group and size class would see the largest relative increase in colony area over one year of linear extension, and applying that growth factor to *H*. Since colony (basal) area scales with the square of colony diameter, the relative area growth of a colony at diameter 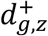 extending by *γ* is the dimensionless ratio 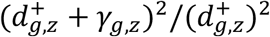; taking the largest such ratio guarantees *C*^∗^ is a sufficiently generous bound that 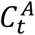 never exceeds *H*.

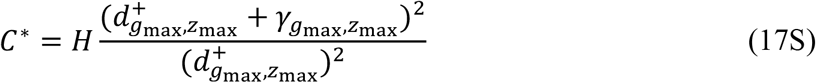

where 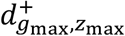 and 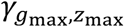 are the upper bound and linear extension of the functional group *g*_max_ and size class *z*_max_ that jointly maximize 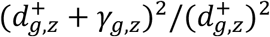, such that 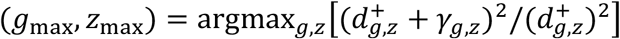, taken over all functional groups *g* and non-terminal size classes *z* = 1, …, *Z* − 1.

One can see that if 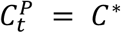, we have 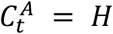. For all other values of 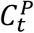, we have 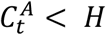, which means the adjusted projected cover will never outgrow the habitable area.

#### 7.4.5 Spatial competition coefficient

As mentioned, the spatial competition scale factor is such that, if we use Eq. (15S) replacing *γ*_*τσ*_ by *γ*_*τσ*_*κ*_*t*_, it yields 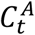:

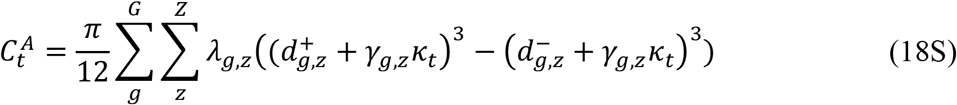

If we solve Eq. (18S) with respect to *κ*_*t*_, we have:

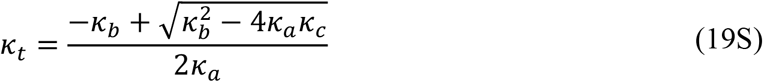

With:

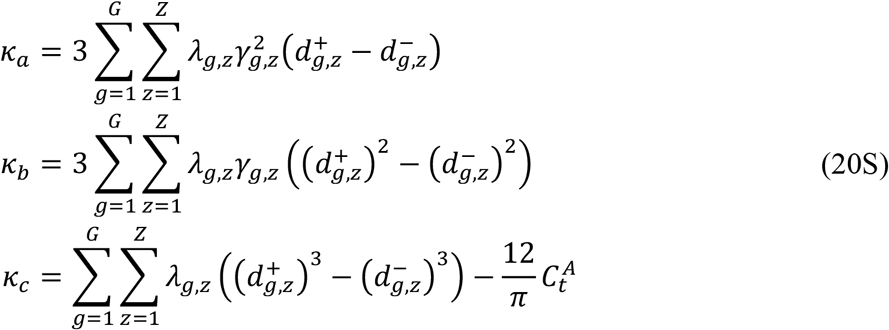

### 7.5 Model calibration and testing

The model was fit to match total coral cover manta tow observations and coral composition observations from the Long-Term Monitoring Program (LTMP) using the Adaptive Differential Evolution algorithm (73), implemented in the BlackBoxOptim.jl Julia package (74). The parameter bound settings, data split and objective function are described in more detail in the following sections

#### 7.5.1 Data split and reef groups

A reef grouping derived from the Great Barrier Reef Marine Park Authority’s bioregions (59, 60). Bioregions were grouped together based on proximity to each other such that each new group had sufficient Long Term Monitoring Program Data for calibration and testing of a coral ecology model.

Data used to calibrate and validate the model is grouped by locations and split into a calibration and a test sets randomly. For 3806 reefs being modelled, only 99 were deemed to have sufficient data for use (4 or more manta tow observations over 10 or more years). Locations with observational data were split within reef groups, ensuring that for each reef group 75% of locations were in the calibration set and 25% were in the test set. This partition was done to ensure that both the calibration and test sets were representative of reef groups. Furthermore, in the case of reef groups with less than eight locations with observation data, the split was guaranteed to assign two locations to the test set. This resulted in 73 calibration locations and 26 test locations.

**Figure S1.**
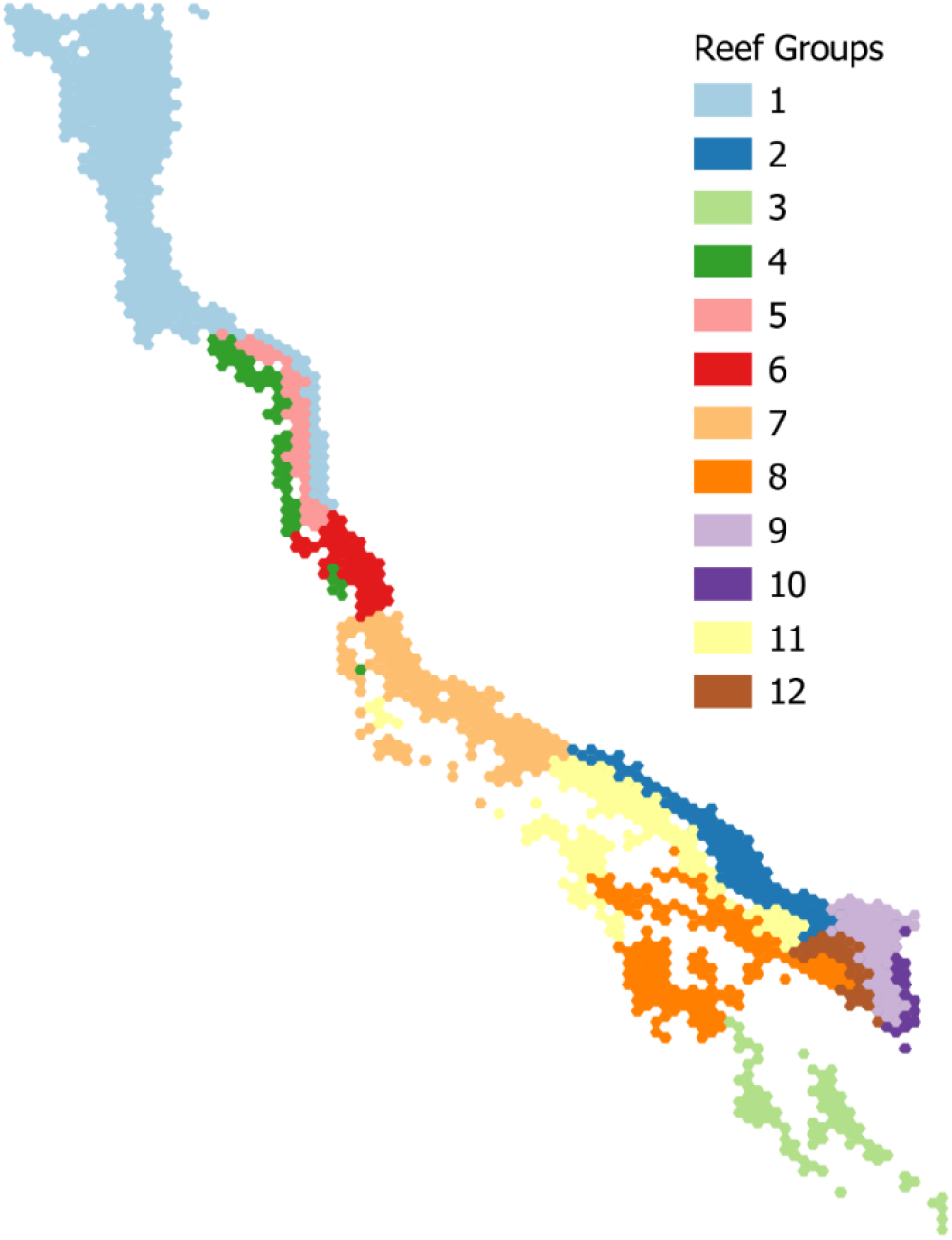
Each of the 12 reef groups used in the calibration.

#### 7.5.2 Objective function

The objective function used to optimize the parameterization of CoralBlox has two components: a total cover error and a coral composition error. AIMS’ Long Term Monitoring Program (LTMP) Manta Tow data was used as the target reef coral cover levels, and coral composition data derived from AIMS’ LTMP photogrammetry data was used as the target coral composition levels.

The total cover error combines a year-weighted mean absolute error over the calibration period with two further terms scoring how well each reef’s observed peak and trough in coral cover are reproduced. The per-year error, calculated across all reefs *l* with an observed value at year *t* (*L*_*t*_, with *N*_*t*_ = |*L*_*t*_| reefs), is given by:

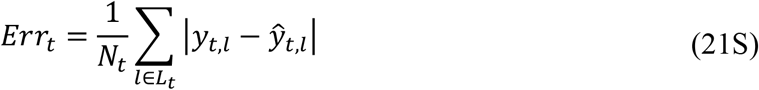

Because CoralBlox is run forward from its initial cover without re-anchoring to observations, error compounds over the course of a simulation. Later years, and years with more reporting reefs, are therefore weighted more heavily when combining the per-year errors into a single reef performance error:

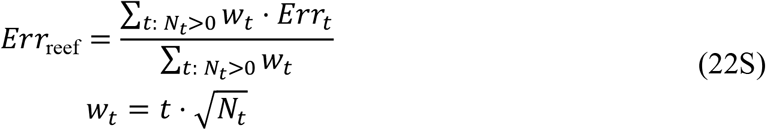

where *t* is the year index (1 for the first calibration year, increasing thereafter), and reefs surveyed in the same year are assumed not to fail independently of one another, so *N*_*t*_ enters as 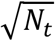 rather than linearly. Years with no reporting reefs are excluded from Eq. (22S).

Modelled coral composition data derived from LTMP photogrammetry data was used to inform the initial coral composition at specific locations and as the target coral composition for subsequent historic years. Coral composition data was not recorded at all locations where manta tow coral cover levels are recorded; the assumption is that composition at fixed-site photogrammetry locations is representative of composition at nearby manta tow locations. Compositional mismatch between simulated and observed functional group cover was calculated using Bray-Curtis dissimilarity between the observed (*c*_*t,g,l*_) and modelled (ĉ _*t,g,l*_) proportional cover of each functional group *g*, and averaged into a single Coral Composition Error (CCE). Bray-Curtis dissimilarity was used in preference to a correlation-based measure for two reasons: a correlation coefficient computed across only five functional groups is statistically unstable, and functional group proportions are compositional (closure-bound, summing to one), which makes correlation among them prone to spurious signal regardless of sample size. Bray-Curtis compares the two composition vectors directly, without any correlation or sampling estimation step, so neither issue applies:

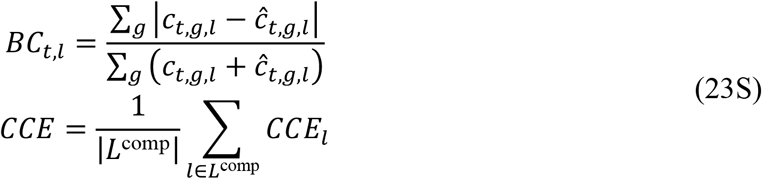

where *CCE*_*l*_ is the mean of *BC*_*t,l*_ over the years at reef *l* where all functional groups’ composition was observed, and *L*^comp^ is the set of reefs with composition data. Reefs with no year of fully observed composition data are assigned a neutral *CCE*_*l*_ of 0.5 rather than being excluded from the mean.

Finally, emphasis is placed on minimizing error at the peak and trough of each reef’s observed coral cover trajectory. For each reef *l*, let 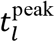 and 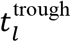 be the years of that reef’s highest and lowest observed coral cover, respectively; the error at these points is calculated using mean absolute error across all reefs *L* with at least one observation:

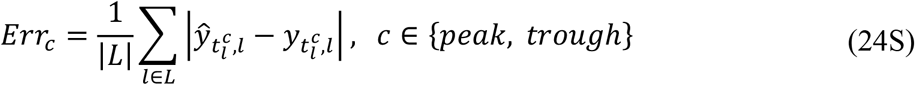

The objective function is then given as

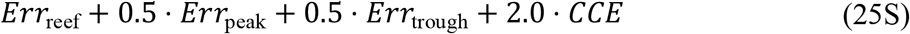

Candidate parameter sets that violate parameter-bound constraints or produce an invalid model run (e.g. a negative recruitment scale factor) are assigned a large fixed penalty instead of being scored by the equations above.

#### 7.5.3 Disturbances datasets

To account for heat stress mortality, we used a historical DHW dataset, derived from observed maximum Degree Heating Weeks (DHWs) data provided by NOAA’s CoralTempv3.1 product (80, 81) for each year and reef (Figure S2). To account for cyclone/storm and COTS mortality, we derived a separate mortality dataset from LTMP disturbance records (year and type of disturbance) with coral cover and benthic composition data. Because coral cover grows naturally between surveys, survival was calculated against a counterfactual undisturbed growth projection of the pre-disturbance cover. This background growth rate was estimated once from each reef’s total cover, then split into per functional group mortality rates using benthic composition data. Locations with disturbances flagged as “unknown” were not included in this mortality-rate calculation.

**Figure S2.**
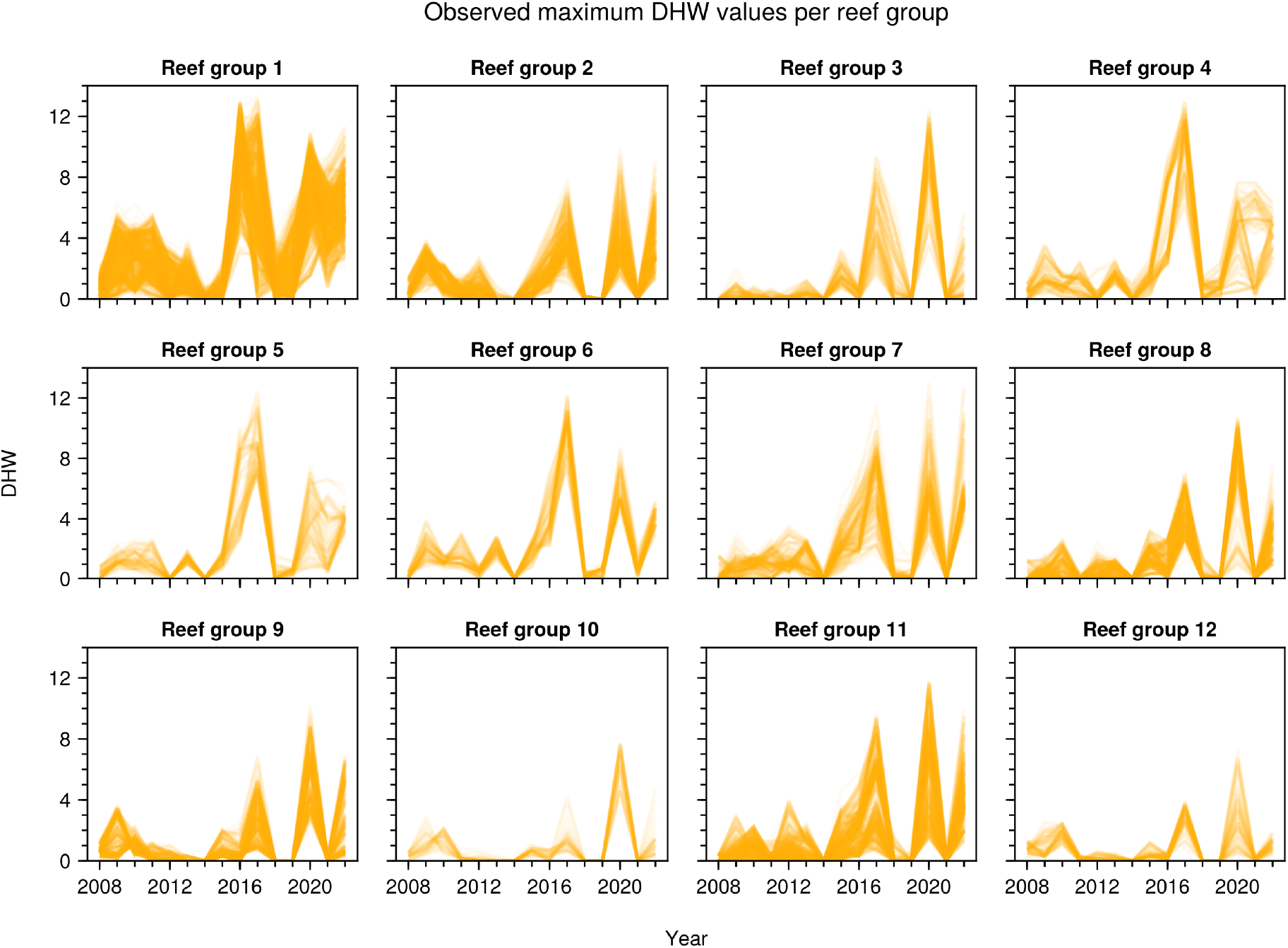
Degree heating weeks (DHW) trajectories for each reef group along the GBR. Individual lines correspond to values at individual reefs within a reef group.

#### 7.5.4 Depth attenuation of DHW

The calibrated *η*^base^ and *η*^mix^ values determine how the effective DHW at depth (Eq. (14)) responds to surface DHW. Figure S3 shows this relationship for a range of depths, using the calibrated parameter values. The depth-refuge effect is strongest at low surface DHW and saturates as surface DHW increases, becoming approximately linear above roughly 10 DHW-weeks.

**Figure S3.**
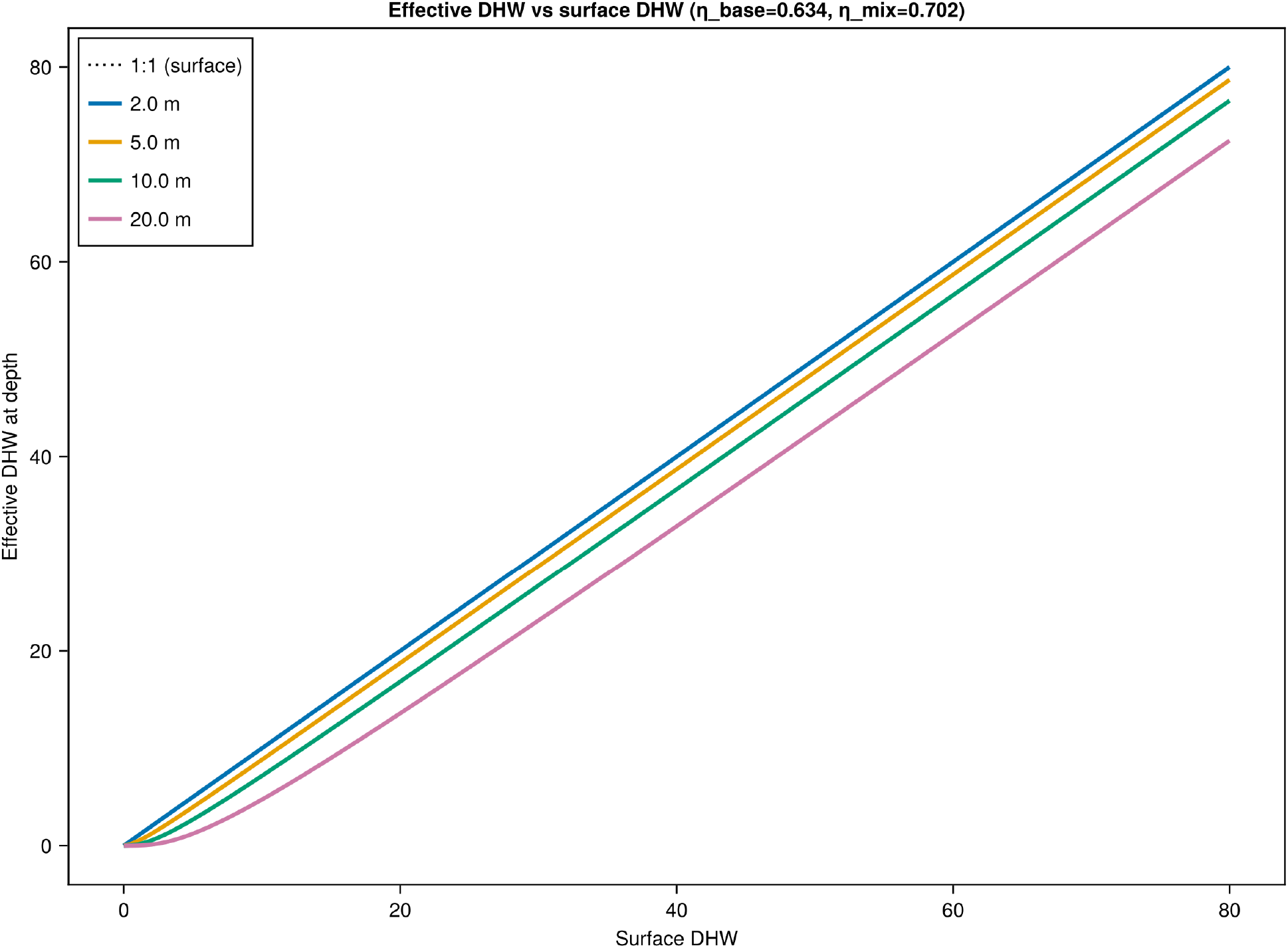
Effective DHW as a function of surface DHW, at four representative depths, using the calibrated *η*^base^ and *η*^mix^ values. The dotted line shows the 1:1 reference (no attenuation, i.e. surface conditions).

### 7.6 Convergence analysis

**Figure S4.**
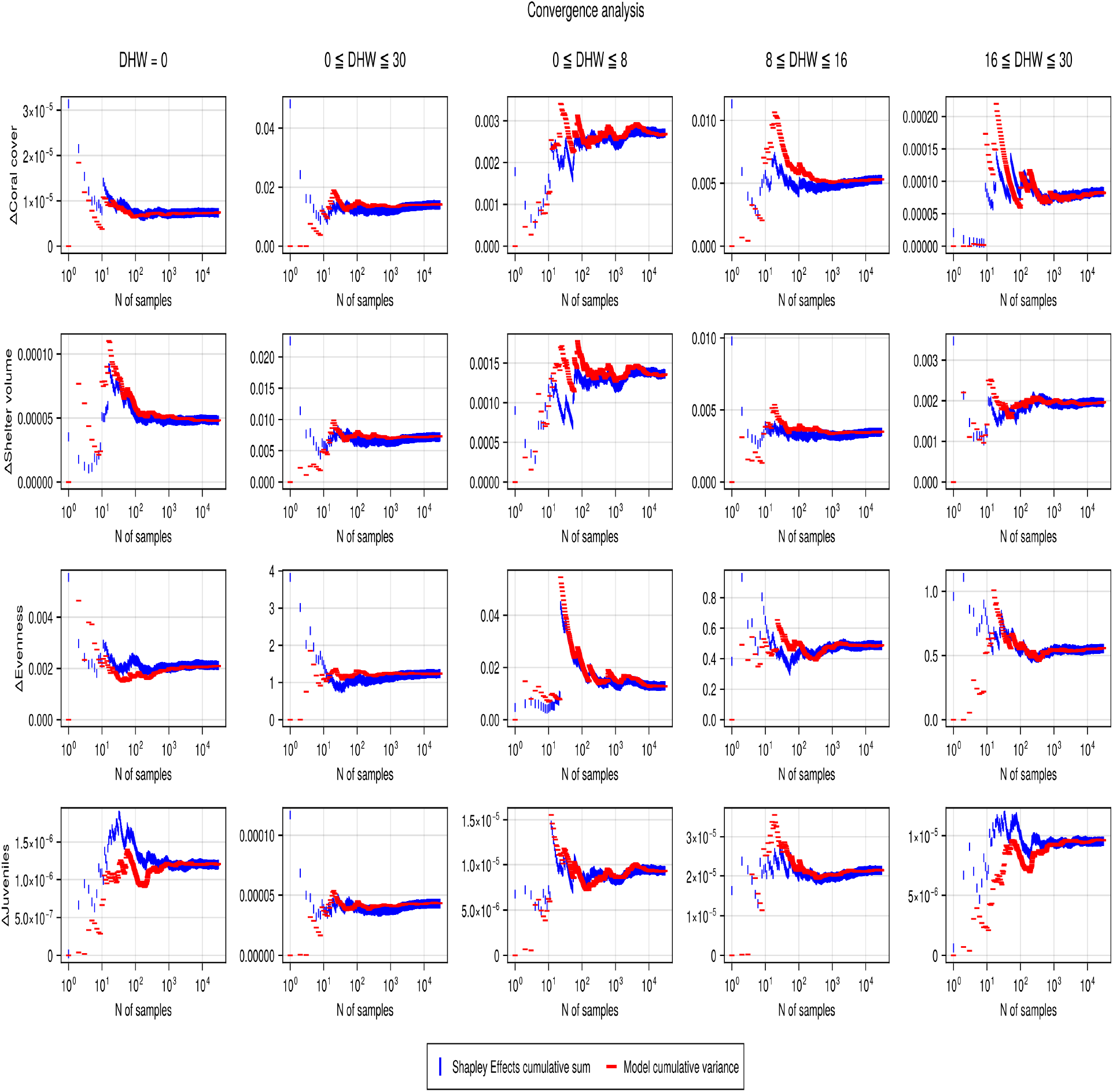
Convergence analysis for each DHW range and metric.

**Table S11.**
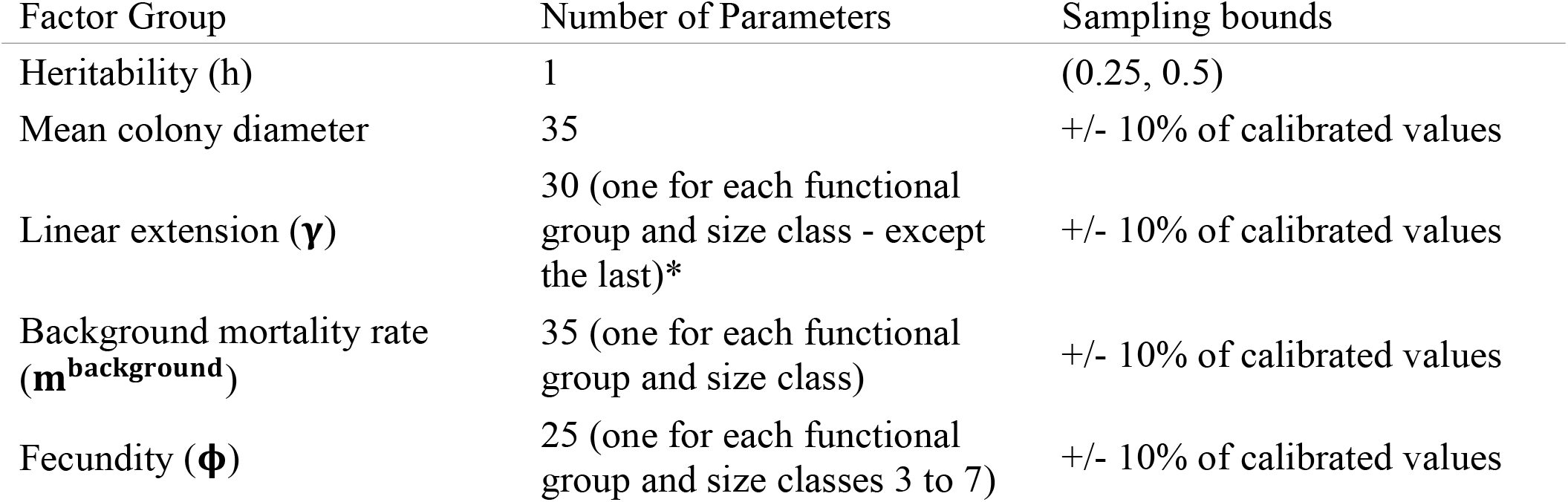
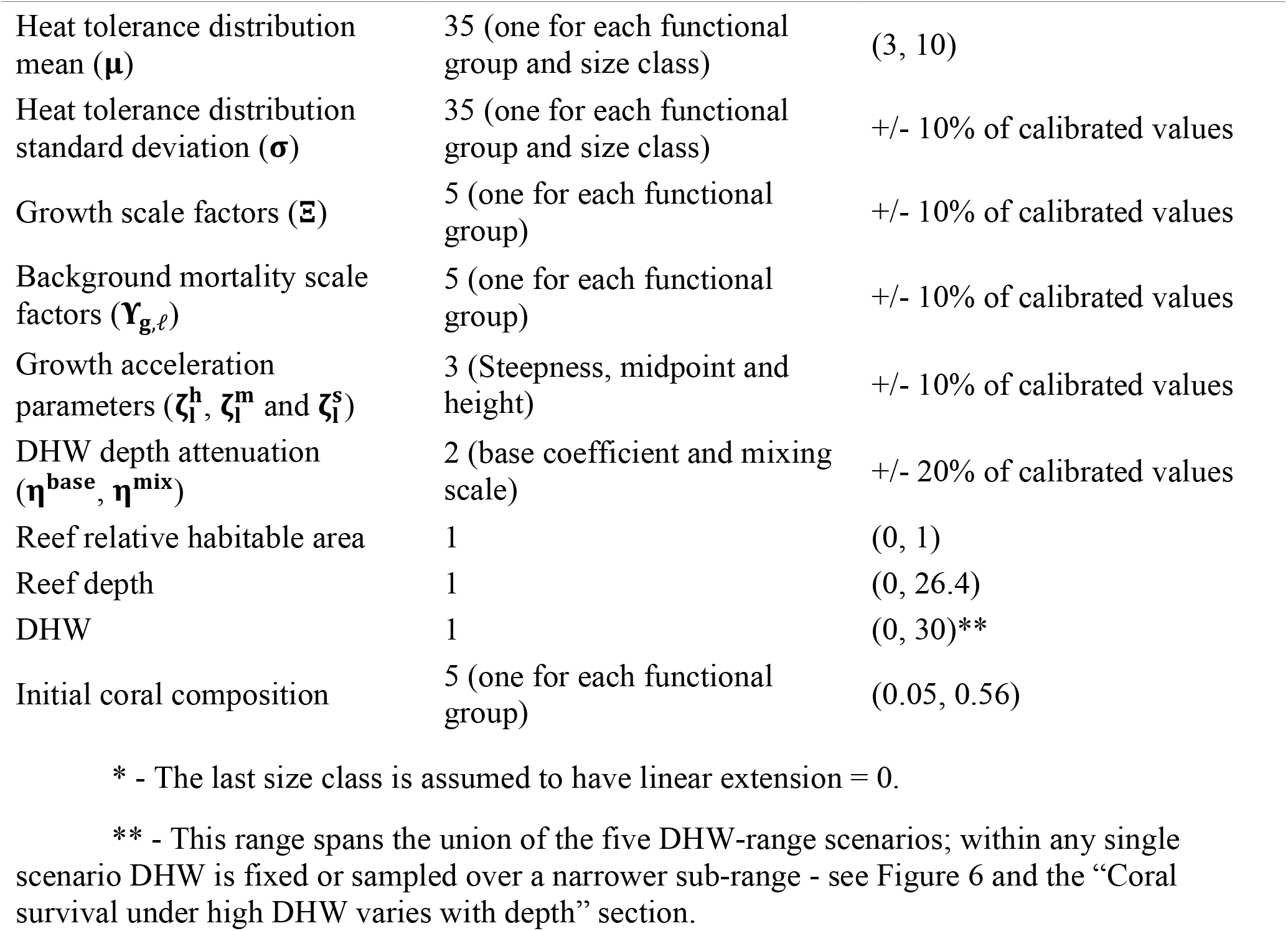
219 factors included in the sensitivity analysis and their respective sampling bounds.

| Factor Group | Number of Parameters | Sampling bounds |
| --- | --- | --- |
| Heritability (h) | 1 | (0.25, 0.5) |
| Mean colony diameter | 35 | +/- 10% of calibrated values |
| Linear extension ($\gamma$) | 30 (one for each functional group and size class - except the last)* | +/- 10% of calibrated values |
| Background mortality rate ($m^{\text{background}}$) | 35 (one for each functional group and size class) | +/- 10% of calibrated values |
| Fecundity ($\phi$) | 25 (one for each functional group and size classes 3 to 7) | +/- 10% of calibrated values |
| Heat tolerance distribution mean ($\mu$) | 35 (one for each functional group and size class) | (3, 10) |
| Heat tolerance distribution standard deviation ($\sigma$) | 35 (one for each functional group and size class) | +/- 10% of calibrated values |
| Growth scale factors ($\Xi$) | 5 (one for each functional group) | +/- 10% of calibrated values |
| Background mortality scale factors ($\Upsilon_{g,\ell}$) | 5 (one for each functional group) | +/- 10% of calibrated values |
| Growth acceleration parameters ($\zeta_l^h$, $\zeta_l^m$ and $\zeta_l^s$) | 3 (Steepness, midpoint and height) | +/- 10% of calibrated values |
| DHW depth attenuation ($\eta^{\text{base}}$, $\eta^{\text{mix}}$) | 2 (base coefficient and mixing scale) | +/- 20% of calibrated values |
| Reef relative habitable area | 1 | (0, 1) |
| Reef depth | 1 | (0, 26.4) |
| DHW | 1 | (0, 30)** |
| Initial coral composition | 5 (one for each functional group) | (0.05, 0.56) |

\* - The last size class is assumed to have linear extension = 0.
\*\* - This range spans the union of the five DHW-range scenarios; within any single scenario DHW is fixed or sampled over a narrower sub-range - see Figure 6 and the “Coral survival under high DHW varies with depth” section.

### 7.7 Per-reef performance metrics

Figures S5-S8 show the full per-reef distributions underlying the aggregate values reported in Table S1: bootstrapped ΔRMSE and SRCC, each with its own 95% confidence interval, for every calibration and test reef, sorted by value. Values and confidence intervals for reefs with 5 or more observed years were computed using a block bootstrap, and for reefs with 4 observed years using an iid bootstrap. We also show the aggregate bootstrapped median with its own 95% confidence interval.

**Figure S5.**
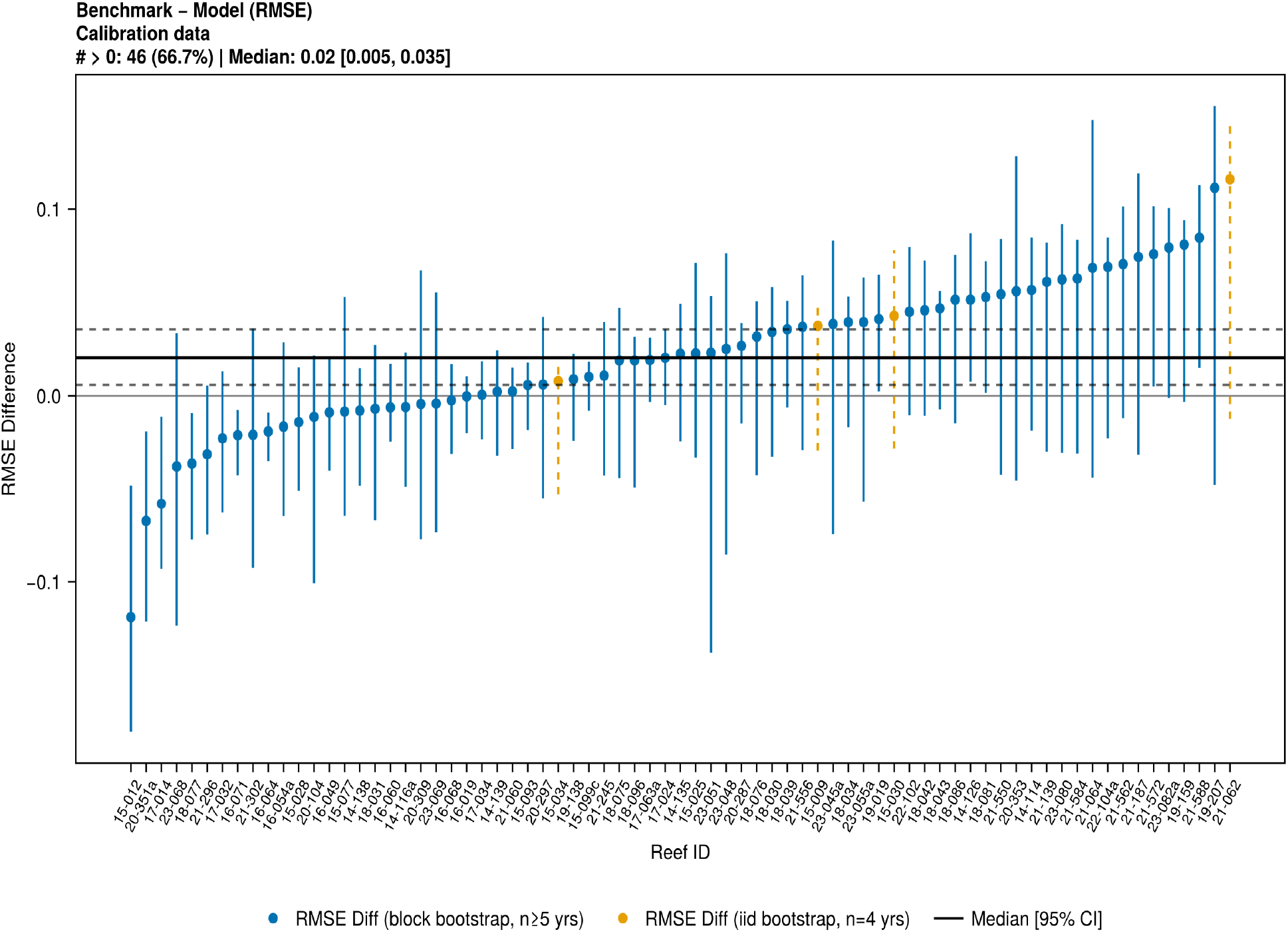
Bootstrapped ΔRMSE (benchmark − model RMSE) for each calibration reef.

**Figure S6.**
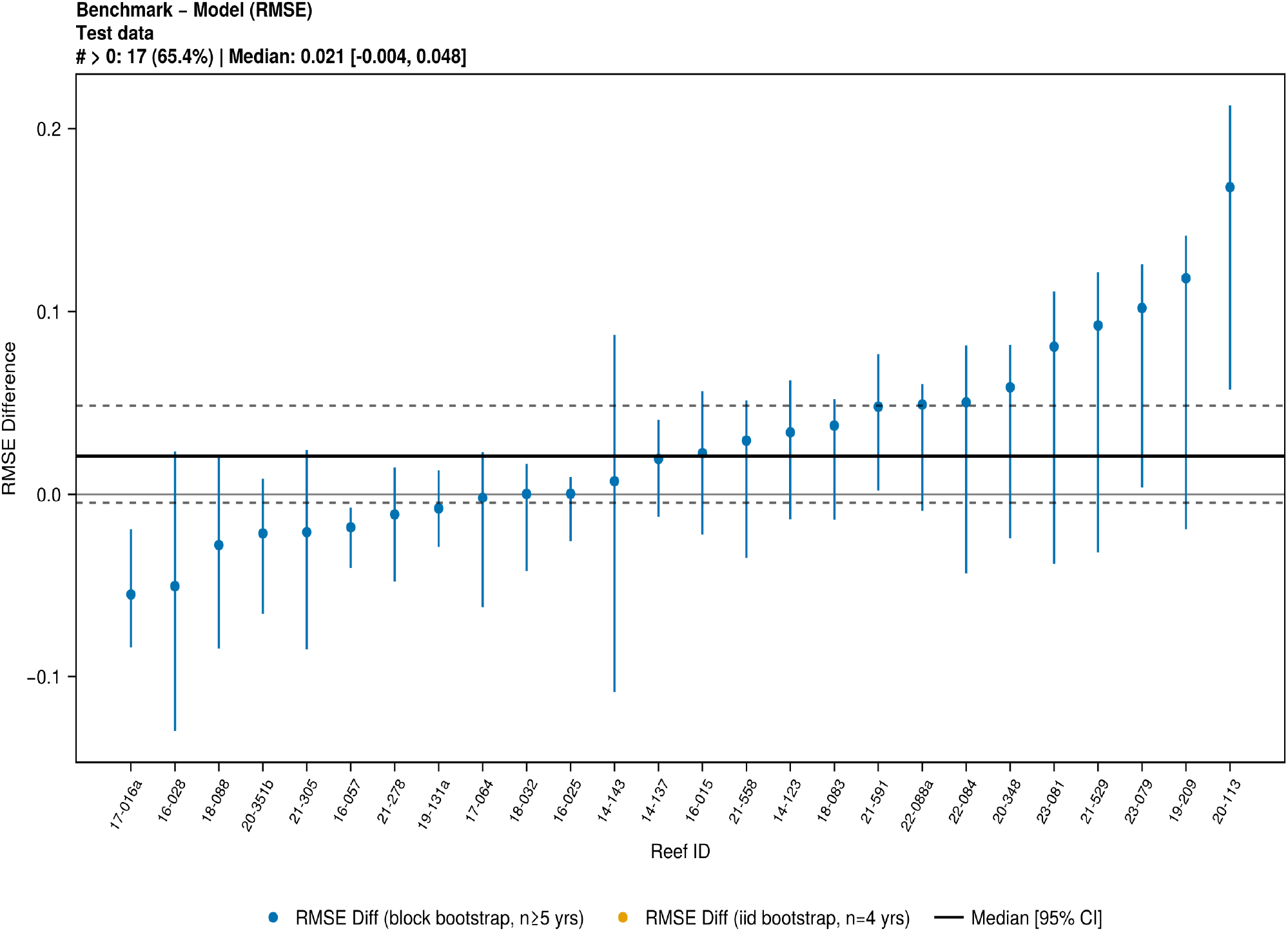
Bootstrapped ΔRMSE (benchmark − model RMSE) for each test reef.

**Figure S7.**
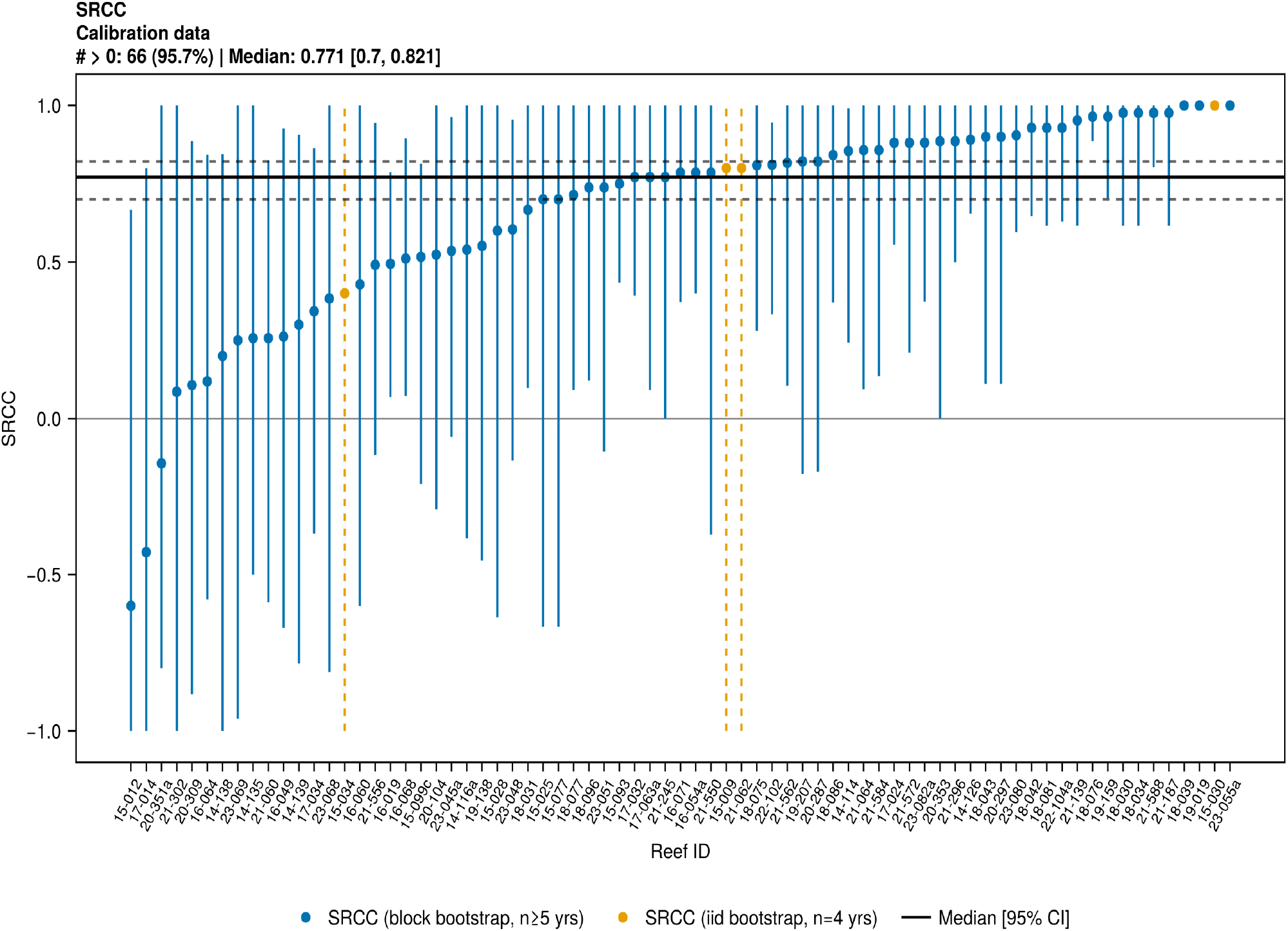
Bootstrapped SRCC for each calibration reef.

**Figure S8.**
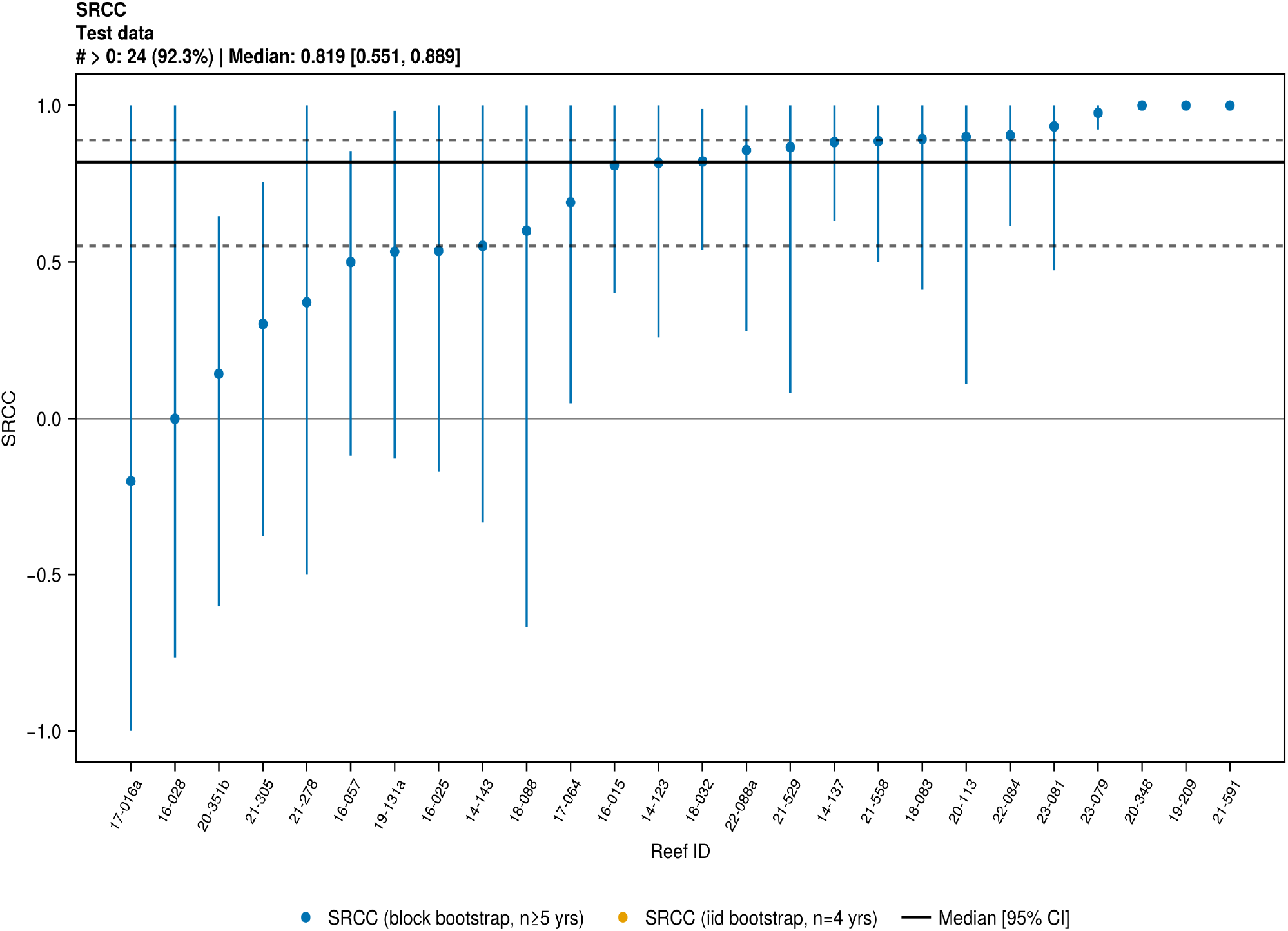
Bootstrapped SRCC for each test reef.

